# mSigHdp: hierarchical Dirichlet process mixture modeling for mutational signature discovery

**DOI:** 10.1101/2022.01.31.478587

**Authors:** Mo Liu, Yang Wu, Nanhai Jiang, Arnoud Boot, Steven G. Rozen

**Author notes:** Denotes equal contribution (Mo Liu, Yang Wu). To whom correspondence may be addressed: SGR.

## Abstract

Mutational signatures are characteristic patterns of mutations caused by endogenous or exogenous mutational processes. These signatures can be discovered by analyzing mutations in large sets of samples – usually somatic mutations in tumor samples. Most programs for discovering mutational signatures are based on non-negative matrix factorization (NMF). Alternatively, signatures can be discovered using hierarchical Dirichlet process (HDP) mixture models, an approach that has been explored less. These models assign mutations to clusters and view each cluster as being generated from the signature of a particular mutational process. Here we describe mSigHdp, an improved approach to using HDP mixture models to discover mutational signatures. We benchmarked mSigHdp and state-of-the-art NMF-based approaches on 4 realistic synthetic data sets. These data sets encompassed 18 cancer types. In total they contained 3.5×10^7^ single-base-substitution mutations representing 32 signatures and 6.1×10^6^ small-insertion-and-deletion mutations representing 13 signatures. For 3 of the 4 data sets, mSigHdp had the best positive predictive value for discovering mutational signatures, and for all 4 data sets, it had the best true positive rate. Its CPU usage was similar to that of the NMF-based approaches. Thus, mSigHdp is an important and practical addition to the set of tools available for discovering mutational signatures.

**Data and code availability:** mSigHdp is available at public repositories https://github.com/steverozen/mSigHdp and https://github.com/steverozen/hdpx. The synthetic data, code for generating the synthetic data, code for running the mutational-signature discovery programs, the main outputs of the programs, and code for analyzing their results and for generating the data figures in this paper are available at https://github.com/Rozen-Lab/mSigHdp sup files. A singularity container with mSigHdp can be downloaded from cloud.sylabs.io with the shell command “singularity pull library://rozen-lab/msighdp/msighdp:2.1.2”. A toy-example Rscript for using this container is at https://github.com/steverozen/mSigHdp/blob/master/data-raw/container_scripts/test_mSigHdp.R.

**Supplementary material:** One excel file of supplementary tables and one PDF file of supplementary figures have been submitted along with this manuscript.

## Introduction

Mutational signature are patterns of mutations that are caused by, and characteristic of, mutational processes or factors (1). Their origins can be endogenous, for example the signatures due to the deamination of 5-methylcytosine or due to activated APOBECs, or exogenous, for example the signatures caused by tobacco smoking or by ultraviolet radiation. The concept of mutational signatures can be applied to various types and classifications of mutations. Single base mutations (single base substitutions, SBSs) are most often classified in the context of the preceding and following bases while considering the mutation to have occurred from a pyrimidine (C or T) (Figure 1A). The mutation classes are, for example, ACA → AAA, ACA → AGA,… CCA → CAA,…, TTT → TGT. Small insertion and deletion mutations (indels) are classified first by their length. Indels of 1 base pair are classified according to the inserted or deleted base and the length of the polynucleotide repeat (if any) in which the indel occurred. Indels of length > 1 base pair are classified according to (i) their length, (ii) whether the indel is in a repeat and the length of the repeat, and, (iii) for deletions, if there is microhomology between the ends of the deleted sequence and the immediately surrounding sequence (Figure 1B). The classification of indels involves several subtleties, detailed at https://www.synapse.org/#!Synapse:syn11801742. For a given biological sample, the counts of mutations in each of the mutational classes constitute the *mutational spectrum* of that sample. Since the samples are usually tumors, for simplicity of exposition, we will refer to biological samples as tumors, understanding that we can discover mutational signatures in other types of biological samples as well.

**Figure 1.**
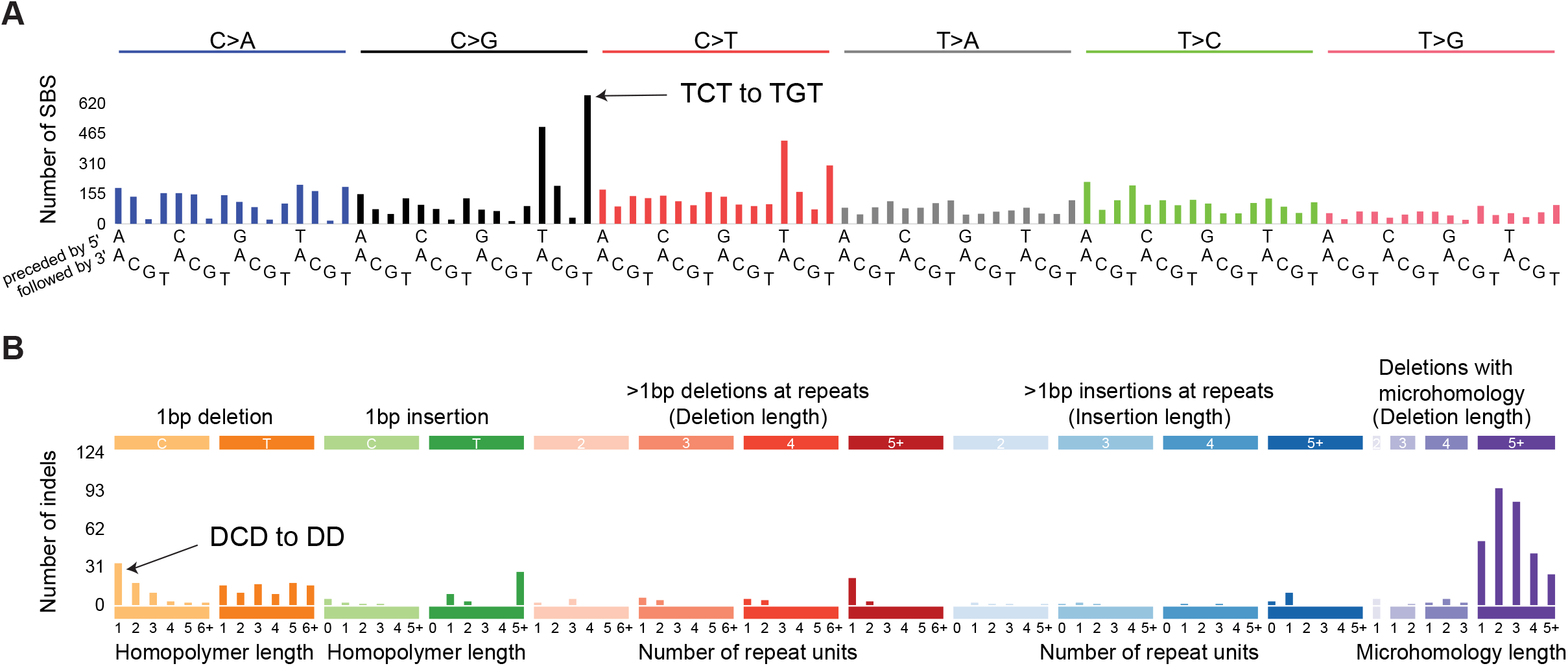
Examples of (A) SBS and (B) indel mutational spectra in a breast cancer. The height of each vertical bar indicates the number of mutations of a particular mutation class, for example, in panel A, 617 mutations from TCT to TGT and in panel B, 34 mutations from DCD to DD (deletion of one C in the middle of two non-C nucleotides, denoted D). This is Breast–AdenoCA::SP116341 from Alexandrov et al. (1) and ICGC/TCGA Pan-Cancer Analysis of Whole Genomes Consortium (36).

For our purposes, a *mutational signature* consists of the expected proportions of mutations in each of the mutation classes, as generated by a particular mutational process. We treat a mutational signature as a multinomial probability vector that maps mutation classes to their probabilities. We sometimes also call these “mutational signature profiles” when we want to emphasize that they are multinomial probability vectors. In our model, the mutational spectrum of 1 tumor is the mutation-class-wise sum of mutations caused by several mutational processes (Figure 2). In this model, a mutational process generates individual mutations by sampling from the multinomial distribution parameterized by the process’s signature, and the spectrum consists of the sum of mutations in each mutational class generated by all the mutational processes operating in the tumor. The task of mutational signature discovery is to infer a set of signatures that can reasonably reconstruct the spectra of a set of tumors. To be useful, the signatures should approximately correspond to underlying biological processes.

**Figure 2.**
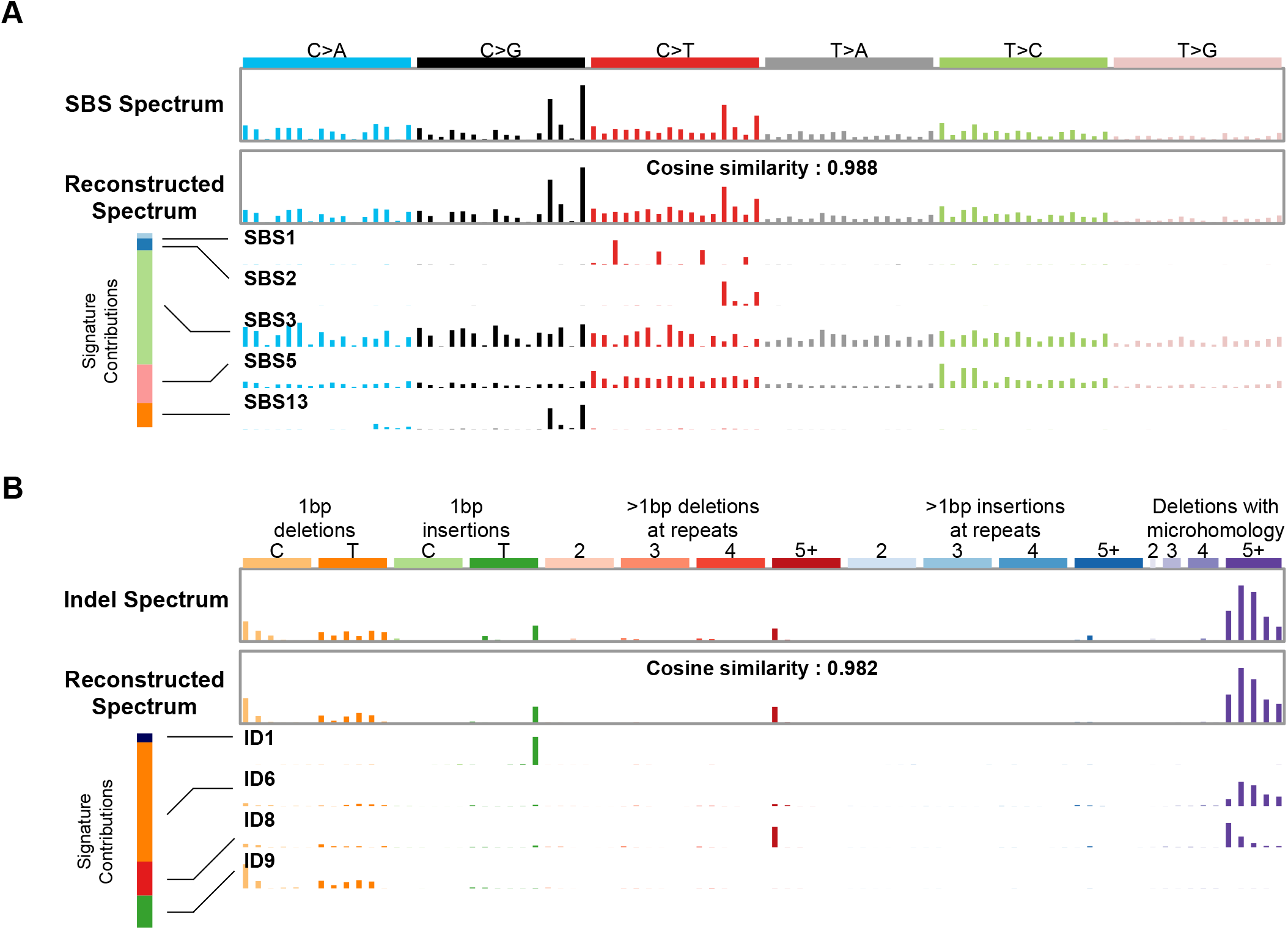
Mutational spectra can be explained as a linear combination of mutations generated by mutational signatures. **(A)** Example of an SBS spectrum that can be reconstructed with a cosine similarity of 0.988 from 5 signatures. The bar at the left shows that mutational signature SBS3 contributed the most mutations to this spectrum. This signature is due to defective homologous-recombination-based DNA repair. The contributions of signatures SBS2 and SBS13 (both due to the activity of endogenous APOBEC cytidine deaminases) are visible in the original and reconstructed spectra. **(B)** The indel spectrum from the same tumor can be reconstructed with a cosine similarity of 0.982 from 4 signatures. The bar at the left shows that mutational signature ID6 contributes the most mutations to this spectrum. Like the SBS signature SBS3, this indel signature is also due to defective homologous-recombination-based DNA repair. Spectra from Breast-AdenoCA::SP116341 (1,36), plotting code in https://github.com/Rozen-Lab/mSigHdp_sup_files.

Early work on discovering mutational signatures relied on approaches based on non-negative matrix factorization (NMF) (2,3) and many subsequent efforts also used NMF-based approaches. To discover mutational signatures, NMF views the tumor spectra as constituting a matrix, with rows being the mutation classes (e.g., for SBSs, CCA → CAA), with the columns being tumors, and with each cell containing the number of mutations of one mutation class for one tumor. NMF then derives an approximate factorization in which the matrix of observed mutation counts is the product of two matrices. The first is a matrix of *K* signatures in which each signature is a column. The second is a matrix in which each element is the number of mutations due to a particular signature in a particular tumor, which we call the *exposure* of that signature in that tumor. Alexandrov and colleagues developed a series of implementations of this concept (1–6). The most recent incarnation of this concept is SigProfilerExtractor, which was assessed along with 13 other approaches in a benchmarking study on synthetic SBS data (6). Signatures discovered using the version in (1) are the basis for the widely used reference set of mutational signatures on the COSMIC website (https://cancer.sanger.ac.uk/signatures/).

As is common in unsupervised learning, a key challenge in mutational signature discovery is determining the number of items, in this case, signatures, that might be present in a set of tumors. NMF-based approaches to mutational signature discovery have deployed several automated or semi-automated approaches to this challenge. SigProfilerExtractor carries out NMF on multiple bootstrapped replicates of the input matrix over a pre-defined range of *K* (the number of signatures to discover). By default, it selects *K* based on a combination of (i) a silhouette analysis of the clustering of signatures discovered across the bootstrap replicates and (ii) the reconstruction errors – how well discovered signatures can reconstruct the input spectra.

In addition to NMF-based approaches, a few approaches to mutational signature discovery have been inspired by probabilistic topic modeling in document classification (7–9). Probabilistic topic modeling is isomorphic to mutational signature discovery: documents correspond to tumors, topics correspond to mutational signatures, and occurrences of individual words in a text correspond to individual mutations in a tumor. In these abstract generative models, topics generate words in characteristic proportions just as mutational signatures generate mutations of different classes in characteristic proportions. Several approaches to probabilistic topic modeling require specification of the number of topics to infer, but hierarchical Dirichlet process (HDP) mixture models estimate the number of topics (or mutational signatures) automatically. Teh et al. (10) showed how to use HDP models for Bayesian estimation of topics and their associated words, and they implemented this approach in C and MATLAB. Roberts (7) showed how to apply this approach to mutational signature discovery, re-implemented the MATLAB portion in R (11), and then applied this implementation to several mutational-signature-discovery tasks (12,13).

Unlike mutational signature discovery by NMF, signature discovery by HDP mixture models focuses on individual mutations. Mutations are assigned to clusters, and a cluster contains a set of mutations that were generated by one mutational process. The classes of the generated mutations in the cluster depend on multinomial draws from the mutational signature of the process. In Teh and colleagues’ implementation, Gibbs sampling is used to estimate the posterior distributions of (i) the number of mutation clusters and (ii) the assignment of observed mutations to the clusters. Clusters of mutations arising from a single mutational process occur in multiple Gibbs samples in a sampling chain. In addition, a single Gibbs sample can contain > 1 cluster of mutations stemming from a single mutational process.

We used the C code from Teh et al. (10) and built upon the R code from Roberts (7),(11) by (i) reducing extremely high R memory usage in the representation of one of the prior distributions, (ii) correcting a problem with the timing of R garbage collection in the R / C interface, and (iii) providing built-in parallelization. Beyond these technical improvements, we made the following conceptual improvements:

1. We reduced the number of false positives by adjusting some of the prior distributions, as described below.
2. We added a facility for downsampling mutation counts in more highly mutated spectra, which we show below markedly reduces false positives in the analysis of SBS data and therefore improves positive predictive values.
3. We implemented a new approach to grouping mutation clusters with similar signature profiles across and within Gibbs samples. This approach avoids grouping mutation clusters with dissimilar signature profiles. This approach stemmed from a more explicit conceptualization of the Gibbs samples as capturing the posterior distribution of the number of mutational processes (i.e., the number of mutational signatures) in addition to capturing the distribution of the proportions of mutations in each mutation class in the process’s signature profile.

Here we describe this implementation of HDP mixture models for mutational signature discovery, termed mSigHdp, and describe the results of benchmarking mSigHdp and 4 additional programs: the original hdp implementation by Roberts and Teh (11), SigProfilerExtractor, SignatureAnalyzer, and signeR, on synthetic SBS and indel mutation data (6,14–17). The additional programs other than the original hdp were those with the best F1 scores in Islam et al. (6).

## Materials and Methods

### Synthetic data generation

We generated 2 sets of synthetic SBS and 2 sets of synthetic indel mutation data, denoted SBS_set1, SBS_set2, indel_set1, and indel_set2. We based these data on (i) the signature profiles from COSMIC (v3.2, https://cog.sanger.ac.uk/cosmic-signatures-production/documents/COSMIC_v3.2_SBS_GRCh37.txt, https://cog.sanger.ac.uk/cosmic-signatures-production/documents/COSMIC_v3.2_ID_GRCh37.txt) and (ii) the numbers of mutations due to each signature in each tumor (the exposure matrix) as estimated by Alexandrov et al. (1). The rows of an exposure matrix are signatures, and the columns are tumors. Each cell contains the number of mutations caused by a particular signature in a particular tumor.

For each cancer type, we took parameters for the synthetic data from the corresponding exposure matrix in Alexandrov et al. (1) as follows:

Let *p_s_* be the probability of occurrence of a signature, *s*, in a tumor of that cancer type. We estimate this from the proportion of tumors of that type in the exposure matrix that have the signature (i.e. have mutation counts > 0 for *s* in the tumor).

Let *μ_s_* be the expected mean of the number of mutations due to *s* in the exposure matrix for those tumors of the given cancer type that have exposure to *s*.

Let *size_s_* be the negative-binomial dispersion parameter of the number of mutations due to *s*, estimated from the dispersion of the number of mutations due to s in the exposure matrix for those tumors of the given cancer type that have exposure to *s*.

We used the R package *fitdistrplus* (18) to get maximum likelihood estimates of parameters *μ_s_* and *size_s_* in the data in Alexandrov et al. (1).

To generate a synthetic spectrum for one tumor type, we decide whether s is present by a random draw from *Bernoulli*(*p_s_*). If s is present, we draw the number of mutations *m_s_* due to s from *NB*(*μ_s_,size_s_*) where *NB* denotes a negative-binomial distribution. For each signature, *s,* present in the tumor, we multiplied *m_s_* by the signature profile vector to get the numbers of mutations in each mutation class, *m_s, t_*, where *t* is a mutation class (e.g., CCA → CAA). To incorporate realistic resampling noise, for each mutation class, t, and signature, *s,* we drew *m_s,t,d_*, the number of mutations of class *t* due to *s*, from *NB*(*m_s,t_, d*), where *d* is the negative binomial dispersion parameter. We selected d = 30 for SBSs and d = 1Q for indels by selecting values that resulted in spectrum-reconstruction accuracies similar to those seen in real data (Supplementary Figures S1, S2). Then, to construct the full synthetic spectrum, we set the number of mutations for each *t* as ∑_*s*_ *m_s,t,d_* and rounded it to an integer.

The synthetic data sets and the code to generate the synthetic data sets are at https://github.com/Rozen-Lab/mSigHdp_sup_files. Supplementary Tables S1 and S2 summarize the signatures and cancer types represented in the synthetic data.

### Running programs for signature discovery

Code to run the signature discovery programs and analyze their output is at https://github.com/Rozen-Lab/mSigHdp_sup_files. Supplementary Table S3 lists the program versions and arguments used. Supplementary Table S4 summarizes signature discovery results for each run. Supplementary Table S5 summarizes CPU usage for each run for which we report CPU times.

### mSigHdp implementation notes

The package hdpx (https://github.com/steverozen/hdpx/ contains the original C code from Teh et al. (10) and much of the R code from Roberts (7). However, in hdpx, the algorithm for finding sets of mutation clusters with similar signature profiles is completely new. We also added diagnostic plots to help assess the reliability of each discovered signature. The first is a scatterplot of the proportion of mutations due to a particular signature versus the count of mutations due to the signature for each input spectrum (Supplementary Figure S3A). The second is a plot of the signature plus 5 input spectra that have the highest proportions of mutations due to that signature (Supplementary Figure S3B). We describe the other plots below. The package mSigHdp (https://github.com/steverozen/mSigHdp/) provides functions for parallelizing and checkpointing the burnin and Gibbs sampling. Checkpointing allows one to extend the burnin if chains have not reached equilibrium distributions. mSigHdp uses the core R function parallel::mclapply for parallelizing over multiple threads, and because mclapply can only use 1 thread on Microsoft Windows, it is not practical to run mSigHdp on Windows.

### Assessing discovered signatures

We assessed how well the discovered signatures reflected the ground truth signatures in the synthetic mutation data as follows. We first matched all the discovered signatures to the ground truth signatures using the function TP_FP_FN_avg_sim in R package mSigTools (https://github.com/Rozen-Lab/mSigTools). This function first computes cosine distances (1 – cosine similarity) and sets cosine distances > 0.1 to a large value, *L.* It then uses the “Hungarian algorithm” (19) to find a matching between discovered and ground truth signatures that minimizes the total cosine distance. Matches with cosine distance *L* are then discarded. After this, discovered signatures that match a ground truth signature are considered true positive matches, any remaining discovered signatures are considered false positives, and any ground truth signatures with no matching discovered signature are considered false negatives.

## Results

### Improvements to discovering mutational signatures with HDP mixture models compared to previous implementations

Hierarchical Dirichlet process (HDP) mixture modeling and its application to mutational signature discovery are described in (7,10). Figure 3 shows an HDP mixture model of mutations generated from mutational signatures. In this conceptualization, signatures model real-world mutational processes. To generate a mutation in, for example, tumor 1 (lower left), one first selects a mutational signature (representing a mutational process) from tumor 1’s Dirichlet process, DP*. This mutational signature is probably one already known to DP*, but it might be a new signature drawn from DP*’s base distribution, DP_1_. Dirichlet-process concentration parameters (not shown in Figure 3) partly govern whether an existing signature is selected or whether a new signature is drawn from the base distribution. If the signature is drawn from DP_1_, then the signature is also probably already known to DP_1_, but if not, it is drawn from DP_1_’s base distribution, DP_0_. If the signature is drawn from DP_0_, it is once again probably already known to DP_0_, but if not, it is drawn from DP_0_’s base distribution, *H. H* is a uniform distribution of an infinite number of possible signatures. More formally, for the case of SBS mutations in trinucleotide context, in which there are 96 mutation classes, *H* is uniform in a 95-simplex, as specified by a symmetric Dirichlet distribution (not a Dirichlet process) with all concentration parameters = 1 (7,10). Once the signature of the mutation is known, the mutation’s class is selected by a multinomial draw from the mutational signature profile. (Recall that a signature profile is a multinomial probability vector.) The number of possible mutational signatures in *H* is unbounded, but depending on the Dirichlet-process concentration parameters, brand new signatures are rarely drawn from *H*.

**Figure 3.**
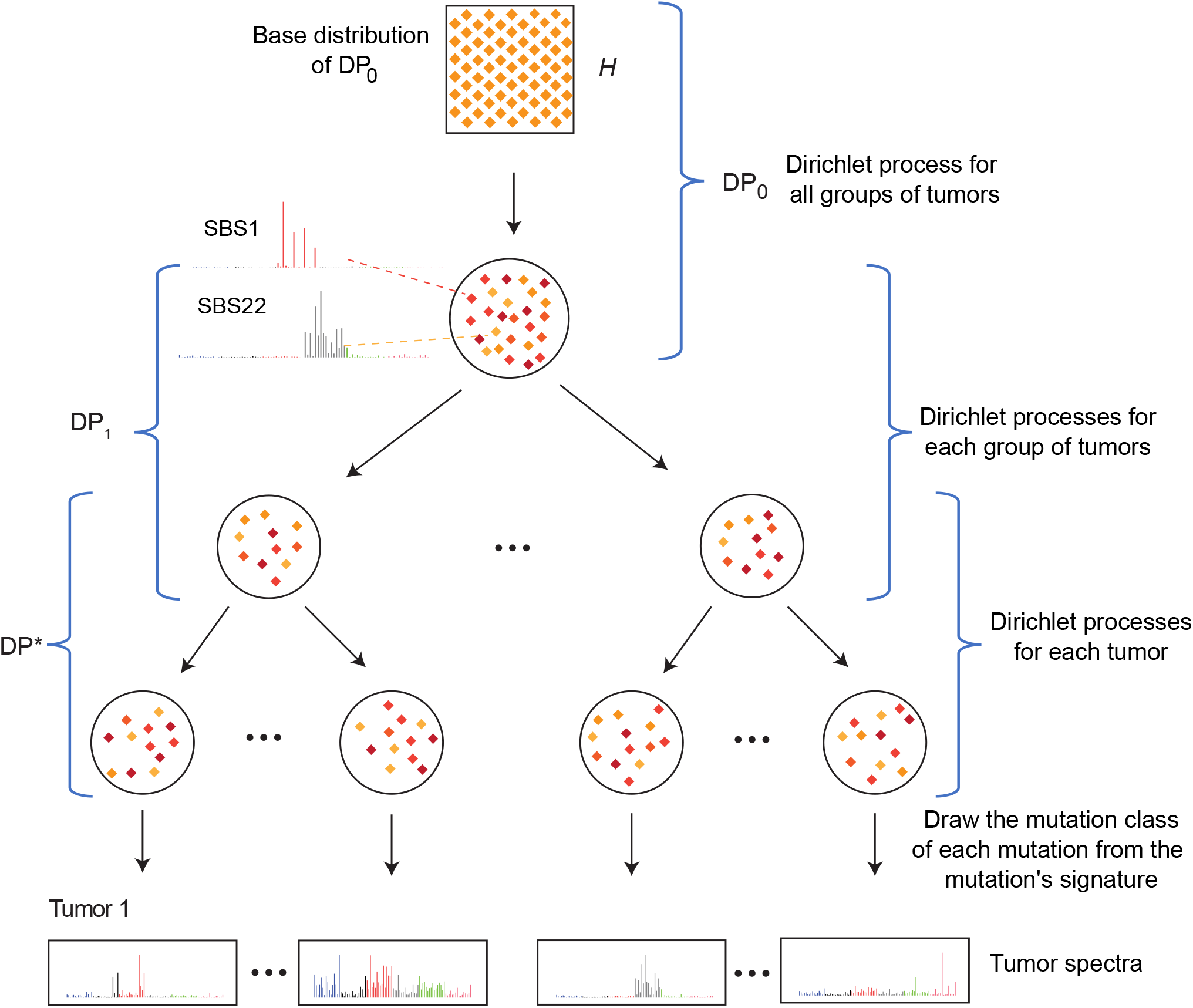
Schematic of a 3-layer HDP mixture model for mutational signature discovery. Please see the text for a full description. Each large circle represents a distribution over signatures in a Dirichlet process. *H* is the base distribution of Dirichlet process *DP_0_, DP_0_* is the base distribution of *DP_1_*, and *DP_1_* is the base distribution of *DP*\*. The small orange diamonds inside each circle represent individual signatures, with darker shades of color indicating a higher probability of that signature in the distribution. Two SBS signatures, SBS1 and SBS22, are shown at DP_0_ as examples. Example observed spectra are shown at the bottom. mSigHdp also supports 2-layer HDP mixture models, in which all tumors have the same parent Dirichlet process, which has *H* as its base distribution.

Teh et al. (10) represented a hierarchical Dirichlet process as a “Chinese restaurant franchise”. The purpose of this representation was to support the implementation of a Bayesian scheme whereby Gibbs sampling could estimate the posterior distribution of the assignments of mutations to clusters generated by a single signature. The Gibbs sampling scheme estimates the posterior distribution of the assignment of customers to tables (which also defines the assignment of mutations to signatures, since each table has a single signature). From the signature, one can also estimate the distribution of the classes of the individual mutations (e.g., CCA → CAA). Each Gibbs sample contains the number of mutations of each mutation class associated with each signature across all tables in all restaurants; here we will call these “mutation clusters”. Teh et al. (10) called these “clusters of category counts”, where “category” corresponds to “mutation class”. In the software, these are represented as matrices in which rows are mutation classes and columns represent different mutation clusters due to different signatures.

An important issue is that across the many Gibbs samples of the posterior distribution, each Gibbs sample represents a signature by its realization as one or more mutation clusters, and across the Gibbs samples, these clusters differ somewhat in their proportions of mutations in each mutation class (Figure 4A). These realizations must be combined to derive point estimates of the signature profiles while noting which signature profiles are common across the Gibbs samples and therefore well supported. The implementation described in Roberts (7) appeared to have high false discovery rates on real data, with a PPV of ~0.5 as estimated from Robert’s Figures 4.13 and D.17. We hypothesized that these high false discovery rates were due to the techniques for combining mutation clusters in (7,11), possibly exacerbated by the use of 1 as the beta parameter of the gamma-distribution prior for the Dirichlet-process concentration parameters. Our subsequent benchmarking confirmed that the previous technique for combining mutation cluster was the major contributor to high false discovery rate, while the use of 1 as the beta parameter of the gamma-distribution prior for the Dirichlet-process concentration parameters was a minor contributor (Figure 5).

**Figure 4.**
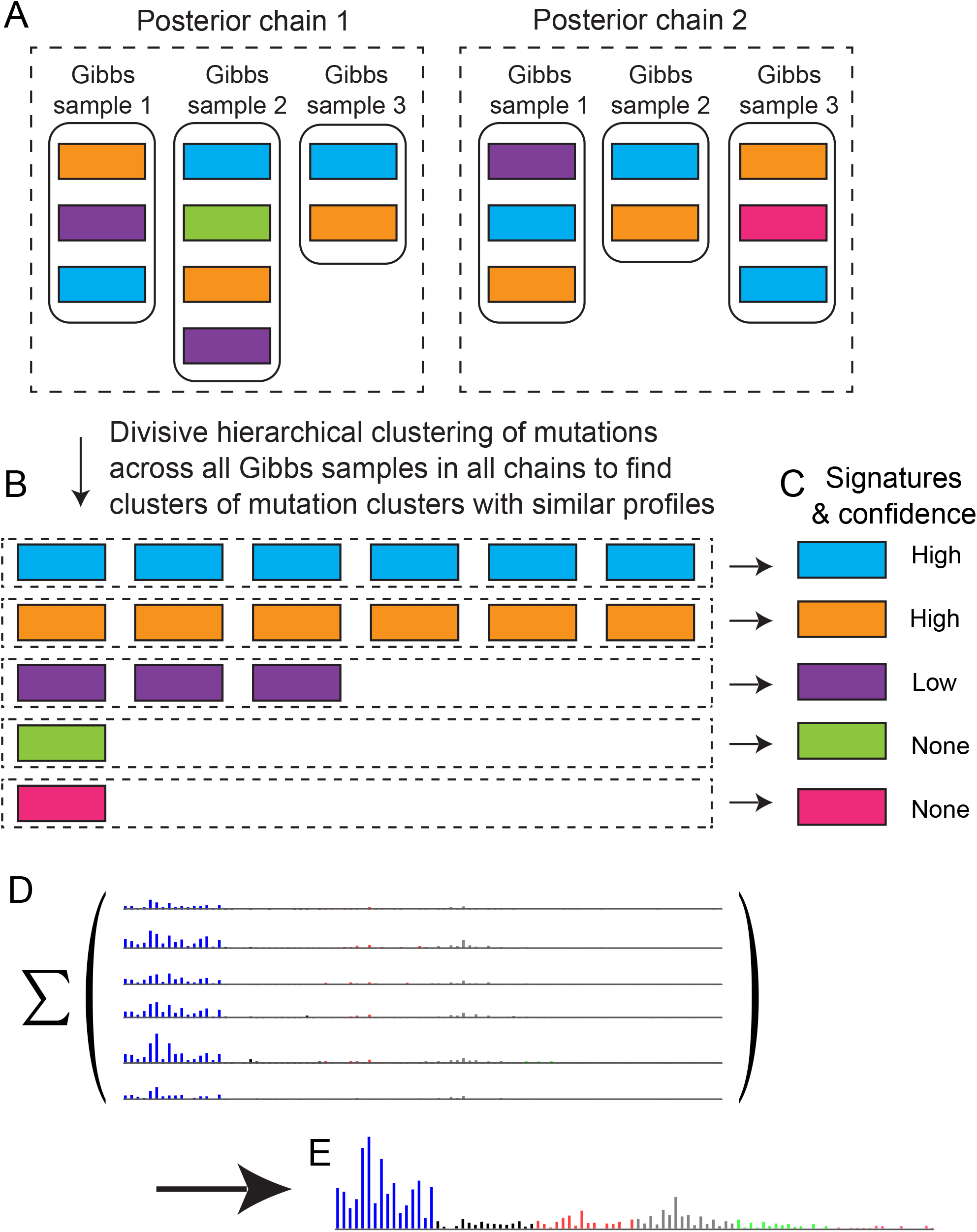
Combining and interpreting mutation clusters collected by Gibbs sampling. **(A)** Gibbs samples from 2 posterior chains in a highly simplified schematic. Each rectangle represents a mutation cluster, and mutation clusters with similar profiles (similar proportions of counts of mutations in each mutation class) have the same color. A typical configuration would be 20 posterior chains in parallel, with 200 Gibbs samples per chain and with 100 iterations between every 2 Gibbs samples. (**B)** Each row (dashed rectangle) is a group of mutation clusters with similar signature profiles (proportions of mutations in each mutation class) as grouped by divisive hierarchical clustering across all Gibbs samples from all sampling chains. We interpret these groups as containing mutation clusters from one signature. **(C)** To infer signatures, in each group of mutation clusters (dashed rectangles in panel B), the counts of mutations of each class are summed, and then these counts are divided by the total number of mutations to yield mutation signatures. **(D, E)** One row of B and C (e.g. the row of blue rectangles) in detail. Each blue rectangle in B is a mutation cluster that is represented by its mutational spectrum in D. The height of each bar in the spectrum indicates the number of mutations of a particular class. Panel E shows the results of summing the mutations of each mutation class over the mutational spectra in D. Dividing by the total number of mutations (not shown) yields the signature profile, corresponding to the blue rectangle in C. In panel C, we have high confidence in signatures that are represented by mutation clusters in all or most of the Gibbs samples, i.e., that are common in the posterior distribution. We suggest a cutoff of ≥ 0.9 of the Gibbs samples as indicating high confidence.

**Figure 5.**
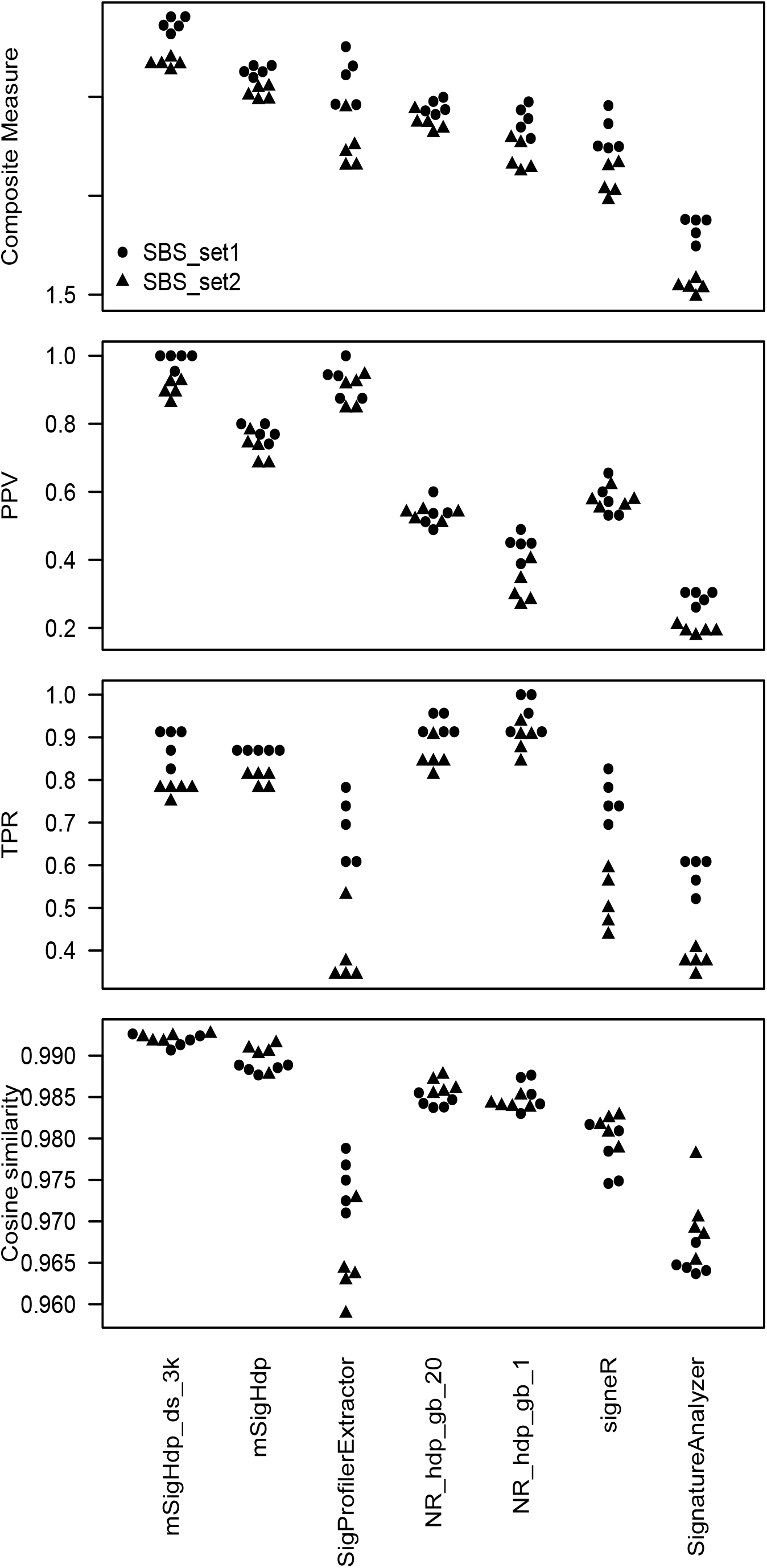
Results on synthetic SBS data. Composite Measure is the sum of PPV (positive predictive value), TPR (true positive rate), and mean cosine similarity. Each circle or triangle represents one measure for one program for one random seed. mSigHdp_ds_3k denotes mSigHdp with a downsampling threshold of 3,000. NR_hdp_gb_20 denotes the implementation in Roberts and Teh (11), with the beta parameter of the gamma-distribution prior for the Dirichlet-process concentration parameters set to 20; NR_hdp_gb_1 is analogous, but with the beta parameter set to 1.

After experimenting with several alternative clustering approaches, we selected divisive hierarchical clustering (20)(function diana in R package cluster, https://cran.r-project.org/package=cluster) to group mutation clusters with similar signature profiles across all Gibbs samples in all chains (dashed-line rectangle in Figure 4B). After this grouping (Figure 4C), we take the signature profile of the mutation-class-wise sums of mutations in all mutation clusters in each group in Figure 4B as the estimate of the underlying mutational signature (Figure 4D,E, Supplementary Figure S4).

In Figure 4C, the “blue” and “orange” signatures are derived from mutation clusters found in all 6 posterior samples in Figure 4A. Because these are samples from the posterior distribution of mutation clusters generated by a mutational process, this indicates strong evidence for the existence of the process generating the “blue” and “orange” mutational signatures. The notion that there is strong evidence for signatures that appear in most posterior samples stems from the theory of Gibbs sampling, in which the Gibbs samples are samples of the posterior distribution of signatures as realized by mutation clusters. Therefore, signatures that appear in most of the posterior samples have strong support, while the others do not. Thus, the “green”and “pink” mutational signatures in panel 4C, which derived from mutation clusters present in only 1 posterior sample, would have negligible support. The “purple” mutational signature in panel 4C, found in only half of the posterior samples, would also have little support. mSigHdp takes an argument that specifies the proportion of posterior samples in which a signature should be found to be considered well supported. This argument defaults to 0.9. Examination of plots that show the Gibbs samples contributing to each signature (Supplementary Figure S5) suggests that false positives would not be further reduced by increasing this argument.

The prior distributions of the Dirichlet-process concentration parameters influence the number of signatures discovered and therefore the false discovery rate (21). The prior distributions are gamma distributions, which have two (hyper)parameters: shape and beta. The location of the gamma distribution decreases as beta increases. The mean of a gamma distribution with shape 1 and beta 1 is nearly 1, and for shape 1 and beta 20 the mean is ~0.05. So compared to a beta of 1, a beta of 20 would encourage smaller concentration parameters, which would make the generation of new signatures from the base distribution less likely. This in turn would shift the posterior distribution of the number of signatures to lower values i.e., fewer signatures. Based on previous analysis (21), for the SBS signature discovery results here, we used beta = 20. For signature discovery in the indel data, we tested several values of beta on the synthetic indel data used here and selected beta = 50 (Supplementary Figure S6). All Dirichlet processes within the mSigHdp models used the same prior for their concentration parameters.

Prompted by the suggestion of a reviewer of a previous version of this manuscript, we tested downsampling as a means to reduce mSigHdp’s computational demands. We found that, for the discovery of SBS mutational signatures, downsampling not only substantially reduced computational requirements, but also substantially improved the false positive rate (i.e. increased the positive predictive value, Figure 5, Table 1, Supplementary Figures S7, function downsample_spectra in package mSigHdp).

**Table 1.** Results on synthetic SBS data. Means and standard deviations (SD) are over the 5 different random seeds. PPV, positive predictive value; TPR, true positive rate. mSigHdp_ds_3k denotes mSigHdp with a downsampling threshold of 3,000. NR_hdp_gb_20 denotes the implementation in (11) Roberts and Teh (11), with the beta parameter of the gamma-distribution prior for the Dirichlet-process concentration parameters set to 20; NR_hdp_gb_1 is analogous with the beta parameter set to 1.

| Data set | Approach | Composite measure |  | Mean PPV | Mean TPR | Mean cosine similarity |
| --- | --- | --- | --- | --- | --- | --- |
|  |  | Mean | SD |  |  |  |
| SBS_set1 | mSigHdp_ds_3k | 2.87 | 0.04 | 0.99 | 0.89 | 0.99 |
|  | mSigHdp | 2.63 | 0.03 | 0.78 | 0.87 | 0.99 |
|  | SigProfilerExtractor | 2.59 | 0.13 | 0.93 | 0.69 | 0.97 |
|  | NR_hdp_gb_20 | 2.45 | 0.04 | 0.54 | 0.93 | 0.98 |
|  | NR_hdp_gb_1 | 2.39 | 0.07 | 0.45 | 0.96 | 0.99 |
|  | signeR | 2.31 | 0.09 | 0.58 | 0.76 | 0.98 |
|  | SignatureAnalyzer | 1.84 | 0.06 | 0.29 | 0.58 | 0.96 |
| SBS_set2 | mSigHdp_ds_3k | 2.67 | 0.02 | 0.90 | 0.78 | 0.99 |
|  | mSigHdp | 2.52 | 0.03 | 0.73 | 0.80 | 0.99 |
|  | SigProfilerExtractor | 2.25 | 0.12 | 0.90 | 0.39 | 0.96 |
|  | NR_hdp_gb_20 | 2.37 | 0.05 | 0.53 | 0.85 | 0.99 |
|  | NR_hdp_gb_1 | 2.20 | 0.08 | 0.32 | 0.89 | 0.98 |
|  | signeR | 2.07 | 0.08 | 0.58 | 0.51 | 0.98 |
|  | SignatureAnalyzer | 1.54 | 0.03 | 0.19 | 0.38 | 0.97 |

### Overall testing strategy

The overall testing strategy was to generate realistic synthetic mutation spectra based on known mutational signatures and then assess the signature discovery approaches on how well they could discover the signatures on which the spectra were based. We generated the synthetic spectra by simulating the operation of known mutational processes, which generated mutations according to their known signature profiles. We call these known signature profiles the “ground-truth signatures”. Within the synthetic data for each cancer type, we modeled the prevalences of the mutational processes on their prevalences in real data and also modeled the numbers of mutations generated by signatures on the distributions of those numbers in real data.

We aimed to include cancer types encompassing as many mutational signatures as reasonably possible, and we included 18 cancer types that were well represented in Alexandrov et al. (1) (Supplementary Tables S1 and S2). Altogether the synthetic spectra contained 32 SBS and 13 indel signatures. The synthetic spectra included both rare and common signatures, with prevalences across all tumor types together that ranged from 0.0019 to 0.97 (SBS_set2) and from 0.012 to 0.95 (indel_set2). There were a total of 1,620 SBS spectra and 3.5×1Q^7^ SBS mutations. There were 3,138 indel spectra and 6.1×1Q^6^ indel mutations.

We evaluated approaches to mutational signature discovery based on (i) the proportion of the ground truth signatures that they discovered, (ii) the proportion of discovered signatures that were not present in the ground truth, and (iii) the similarity between the signature profiles of the discovered and ground truth signatures. In more detail, let *TP* be the number of true-positive signatures, i.e. ground truth signatures that were discovered, let *FP* be the number of false-positive discovered signatures (signatures that were discovered but that were not among the ground truth signatures), and let *FN* be the number of false negatives, i.e. ground truth signatures that were not discovered. We then assessed the performance of a mutational-signature-discovery approach by three measures:

1. Positive predictive value, 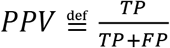
2. True positive rate, 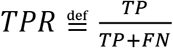
3. The mean of the cosine similarities of true positive signatures to their matching ground truth signatures.

As a summary assessment, we defined the *Composite Measure* as the sum of *PPV, TPR,* and the mean cosine similarity.

### mSigHdp accuracy in discovering SBS signatures

We assessed how well mSigHdp with and without downsampling and 4 other programs discovered SBS mutational signatures on two synthetic data sets (Figure 5, Table 1). Because all the programs that we tested depend on random sampling, their results often vary depending on the initial random seed. Therefore, we ran each program on 5 different seeds, and all programs indeed often generated different results with different starting random seeds.

The Composite Measures of mSigHdp with a downsampling threshold of 3,000 was significantly higher than those of mSigHdp without downsampling and of SigProfilerExtractor (means across all seeds and both synthetic data sets of 2.77 versus 2.57 and 2.42, *p* < 4.4 × 10^-5^ and *p* < 3.3 × 10^-4^, respectively, 2-sided Wilcoxon rank-sum tests, significant at a Bonferroni corrected alpha of 0.025). The Composite Measures of the other approaches were still lower. Details of their results are available in Supplementary Table S4. The mean cosine similarities across true positives were similar across all 4 programs and therefore did not strongly influence the Composite Measures.

Of note, some mSigHdp false positives consisted of essentially a single mutation class (e.g. only TCG → TTG, Supplementary Figure S8). This was also true for mSigHdp without downsampling and for the hdp approach in Roberts and Teh (11). Thus, one should be skeptical of single-mutational-class signatures generated by these approaches from SBS data.

mSigHdp with downsampling had higher TPRs than SigProfilerExtractor. In some cases, SigProfilerExtractor false positive signatures could be decomposed as merges of 2 or 3 false negatives, and considering these as discoveries would increase the assessment of true positives and decrease false negatives (Supplementary Figure S9). However, many false negatives could not be accounted for as members of merged false positives, and this was especially true of most of the false negatives in SBS_set2 (Supplementary Table S4). Discovery of merged signatures is common in mutational-signature analysis. To help address this, SigProfilerExtractor can assess whether discovered signatures are merges of known signatures. While this does not help when discovered signatures are merges of unknown and known signatures, it is useful when discovered signatures are merges of known signatures. Although mSigHdp does not carry out this analysis automatically, it can be done by function find_best_reconstruction_QP in package mSigTools (https://github.com/Rozen-Lab/mSigTools).

Both mSigHdp and SigProfilerExtractor tended to miss less prevalent signatures, but mSigHdp was better able to detect rare signatures (*p* < × 10^-4^ by robust linear regression; see Supplementary Table S7). To investigate this further, we “spiked in” signature SBS35 (which was not discovered by either approach in either SBS data set) at varying concentrations. mSigHdp was able to detect it at lower concentrations than SigProfilerExtractor (*p* < 7 × 10^-3^ by robust linear regression; Supplementary Figure S10).

We also investigated whether SigProfilerExtractor’s true positive rates could be improved by selecting larger values of *K* (the number of signatures extracted, which by default, SigProfilerExtractor selects automatically). At 2 plus the *K* that SigProfilerExtractor had automatically selected (2 + *K* signatures total), it added an average of 1.2 true positives and 0.8 false positives (Supplementary Table S6). At the value of *K* selected by mSigHdp (*K* = 21), SigProfilerExtractor added an average of 2.2 true positives and an average of 1.8 false positives. Thus, SigProfilerExtractor’s true positive rates could indeed be improved by selecting larger values of *K*, albeit at the cost of some increase in the number of false positives. The Composite Measures increased slightly but not significantly at larger values *K* (Supplementary Table S6).

We also studied the extent to which the results of mSigHdp and the other approaches depended on the overall level of resampling noise used in generating the synthetic data (Supplementary Figure S11). For the synthetic data with reduced or no noise, all programs produced better results than when tested on realistic noise. However, the ranking of the programs was substantially unchanged with one exception. This was SignatureAnalyzer, which generated markedly better results for data without resampling noise. Thus, our earlier assessment of SignatureAnalyzer on data without noise (1) was not predictive of its performance on more realistic data.

Finally, as noted above, Figure 5 indicates that the algorithm for grouping raw mutation clusters in the Gibbs samples in (7,11) was the main contributor to a high false discovery rate on real SBS data. This implementation had markedly worse PPVs (^~^0.5) than mSigHdp with downsampling (0.99) and SigProfilerExtractor (0.91). The use of 1 rather than 20 as the beta parameter of the gamma-distribution prior for the Dirichlet-process concentration parameters corresponded to a decrease in PPV from 0.53 to 0.45, and so was a minor contributor.

### mSigHdp accuracy in discovering indel signatures

We assessed how well mSigHdp and the other programs discovered indel mutational signatures in two synthetic data sets (Figure 6, Table 2). Downsampling indel data degraded mSigHdp’s accuracy, and we did not investigate this further (Supplementary Figure S12). mSigHdp had the highest Composite Measure (mean 2.89), and the hdp implementation from Roberts and Teh (11) and Roberts (7) also did well (means 2.80 and 2.78 for two different choices of the prior distribution of the HDP concentration parameter). mSigHdp was significantly better than SigProfilerExtractor (mean Composite Measure 2.89 versus 2.67, *p* < 1.1 × 10^-5^, two-sided Wilcoxon rank-sum test over the results for all random seeds and both synthetic data sets). The other two programs had substantially lower Composite Measures (Table 2, Supplementary Table S4). As with the SBS data, the PPV and TPR were the main contributors to differences in the Composite Measure across the 4 programs.

**Figure 6.**
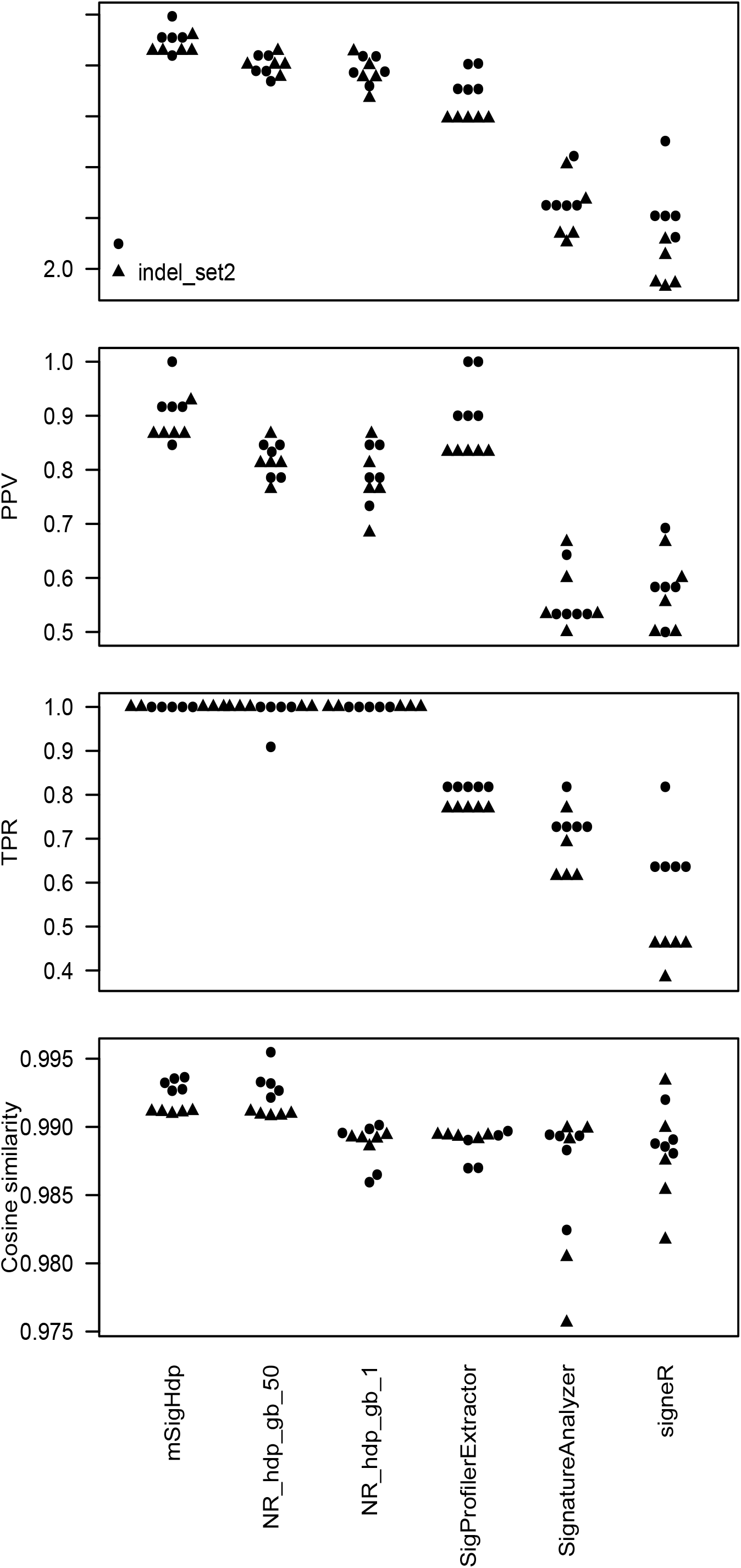
Results on synthetic indel data. Composite Measure is the sum of PPV (positive predictive value), TPR (true positive rate), and mean cosine similarity. Each circle or triangle represents one measure for one program for one random seed. NR_hdp_gb_50 denotes the implementation in Roberts and Teh (11), with the beta parameter of the gamma-distribution prior for the Dirichlet-process concentration parameters set to 50; NR_hdp_gb_1 is analogous, but with the beta parameter set to 1.

**Table 2.**
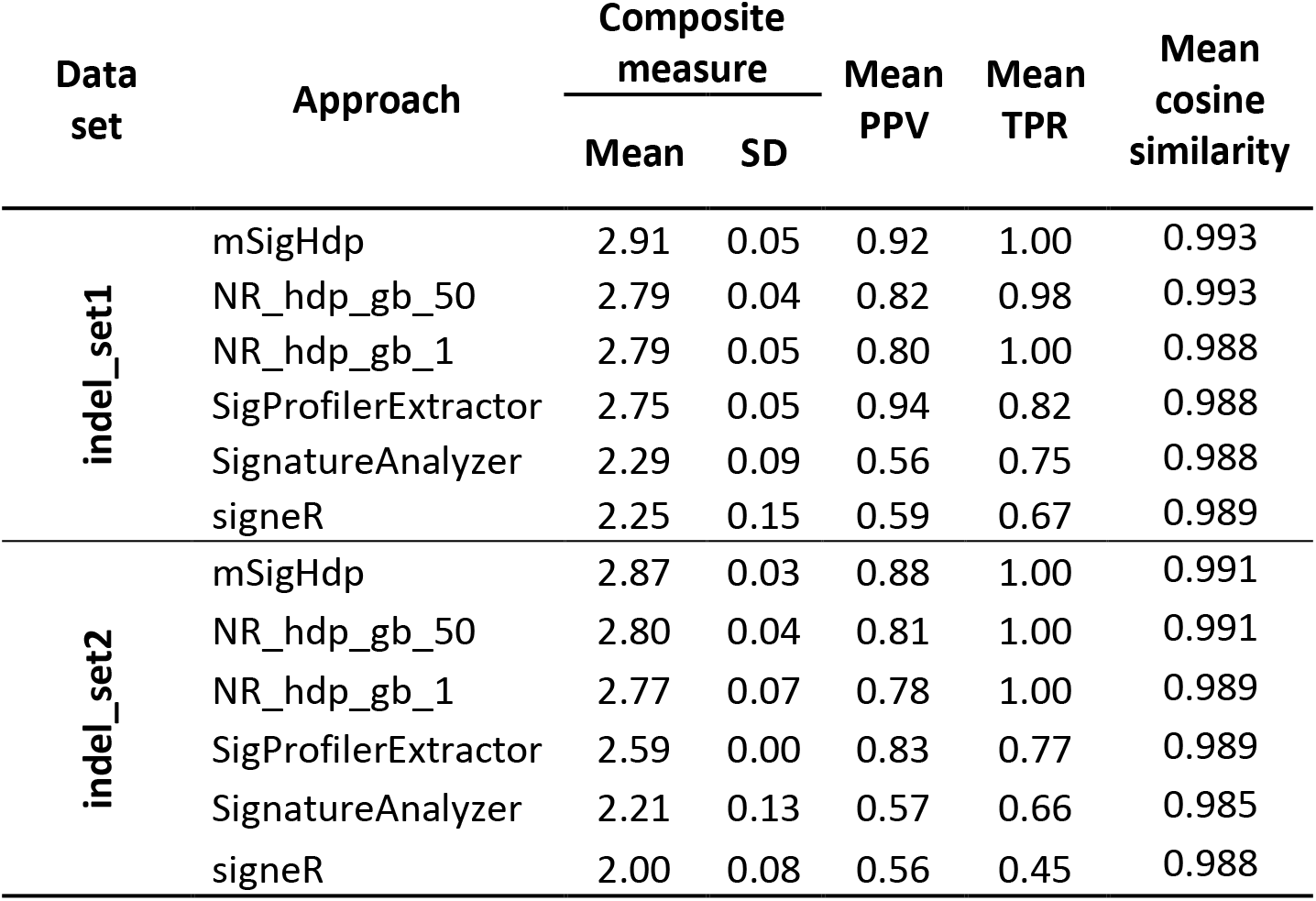
Results on synthetic indel data. Means and standard deviations (SD) are over the 5 different random seeds. PPV, positive predictive value; TPR, true positive rate. NR_hdp_gb_50 is the implementation in Roberts and Teh (11), with the beta parameter of the gamma-distribution prior for the Dirichlet-process concentration parameters set to 50; NR_hdp_gb_1 is analogous with the beta parameter set to 1.

| Data set | Approach | Composite measure |  | Mean PPV | Mean TPR | Mean cosine similarity |
| --- | --- | --- | --- | --- | --- | --- |
|  |  | Mean | SD |  |  |  |
| indel_set1 | mSigHdp | 2.91 | 0.05 | 0.92 | 1.00 | 0.993 |
|  | NR_hdp_gb_50 | 2.79 | 0.04 | 0.82 | 0.98 | 0.993 |
|  | NR_hdp_gb_1 | 2.79 | 0.05 | 0.80 | 1.00 | 0.988 |
|  | SigProfilerExtractor | 2.75 | 0.05 | 0.94 | 0.82 | 0.988 |
|  | SignatureAnalyzer | 2.29 | 0.09 | 0.56 | 0.75 | 0.988 |
|  | signeR | 2.25 | 0.15 | 0.59 | 0.67 | 0.989 |
| indel_set2 | mSigHdp | 2.87 | 0.03 | 0.88 | 1.00 | 0.991 |
|  | NR_hdp_gb_50 | 2.80 | 0.04 | 0.81 | 1.00 | 0.991 |
|  | NR_hdp_gb_1 | 2.77 | 0.07 | 0.78 | 1.00 | 0.989 |
|  | SigProfilerExtractor | 2.59 | 0.00 | 0.83 | 0.77 | 0.989 |
|  | SignatureAnalyzer | 2.21 | 0.13 | 0.57 | 0.66 | 0.985 |
|  | signeR | 2.00 | 0.08 | 0.56 | 0.45 | 0.988 |

mSigHdp and SigProfilerExtractor had similar positive predictive values (0.90 and 0.89). As was the case for SBS data, the HDP implementation from Roberts and Teh (11) had lower positive predictive values for both values of the beta parameter of the gamma-distribution prior of the Dirichlet process hyperparameters (0.82 and 0.79). The main reason again appears to have been the method for combing raw clusters of mutations within and across Gibbs samples.

mSigHdp had a mean of 1.4 false positives across the indel data sets and SigProfilerExtractor had mean of 1.3. mSigHdp discovered a total of 4 different false positives (Supplementary Figure 13), and SigProfilerExtractor discovered a total of 2 different false positives (Supplementary Figure S14). The most common of these was a variant of ID5 in which there were no deletions from TT → T. The other was a singleton that did not resemble any ground-truth signature and that could not be reconstructed from false negatives.

mSigHdp had no false negatives in either indel data set, while other programs had substantial numbers of false negatives. SigProfilerExtractor did not discover ID5 in either indel data set. As noted above, SigProfilerExtractor instead extracted a version of ID lacking TT → T deletions. This was a false positive, as its cosine similarity to ID5 was ~0.8 (Supplementary Figure S14). SigProfilerExtractor also often missed relatively rare signatures ID11 and ID13 in indel_set1 and ID10 and ID12 in indel_set2.

Unlike the SBS signatures, the indel signatures discovered by SigProfilerExtractor at 2 +the selected *K* included no additional true positives but 2 additional false positives (Supplementary Table S6). At *K* as selected by mSigHdp (*K* = 11), there were 0 additional true positives and an average of 1.4 additional false positives. Composite Measures were significantly lower (Supplementary Table S6). Thus, it does not seem that the indel false negatives mainly stemmed from incorrect estimates of *K*.

As we did for SBS data, we investigated the extent to which the results depended on the overall level of resampling noise used in generating the synthetic data (Supplementary Figure S15). Again, the ranking of the programs was substantially unchanged, except for SignatureAnalyzer, which had markedly better results when analyzing data with moderate or no resampling noise.

### CPU usage

For SBS data, mSigHdp with a downsampling threshold parameter of 3,000 required 1.5 times as many CPU resources as SigProfilerExtractor and 9.8 times less than mSigHdp without downsampling (Figure 7, Table 3). CPU usage by mSigHdp with downsampling is dramatically reduced because mSigHdp’s running time is proportional to the number of mutations being analyzed. This in turn is because the time needed for each iteration of the Gibbs sampler depends partly on the number of mutations in the input data set. While mSigHdp required 4.4 times as many CPU resources as SigProfilerExtractor for analyzing the indel data, the overall CPU usage was low, at ^~^4.7 CPU days over 20 threads, amounting to < 6 CPU hours per thread.

**Figure 7.**
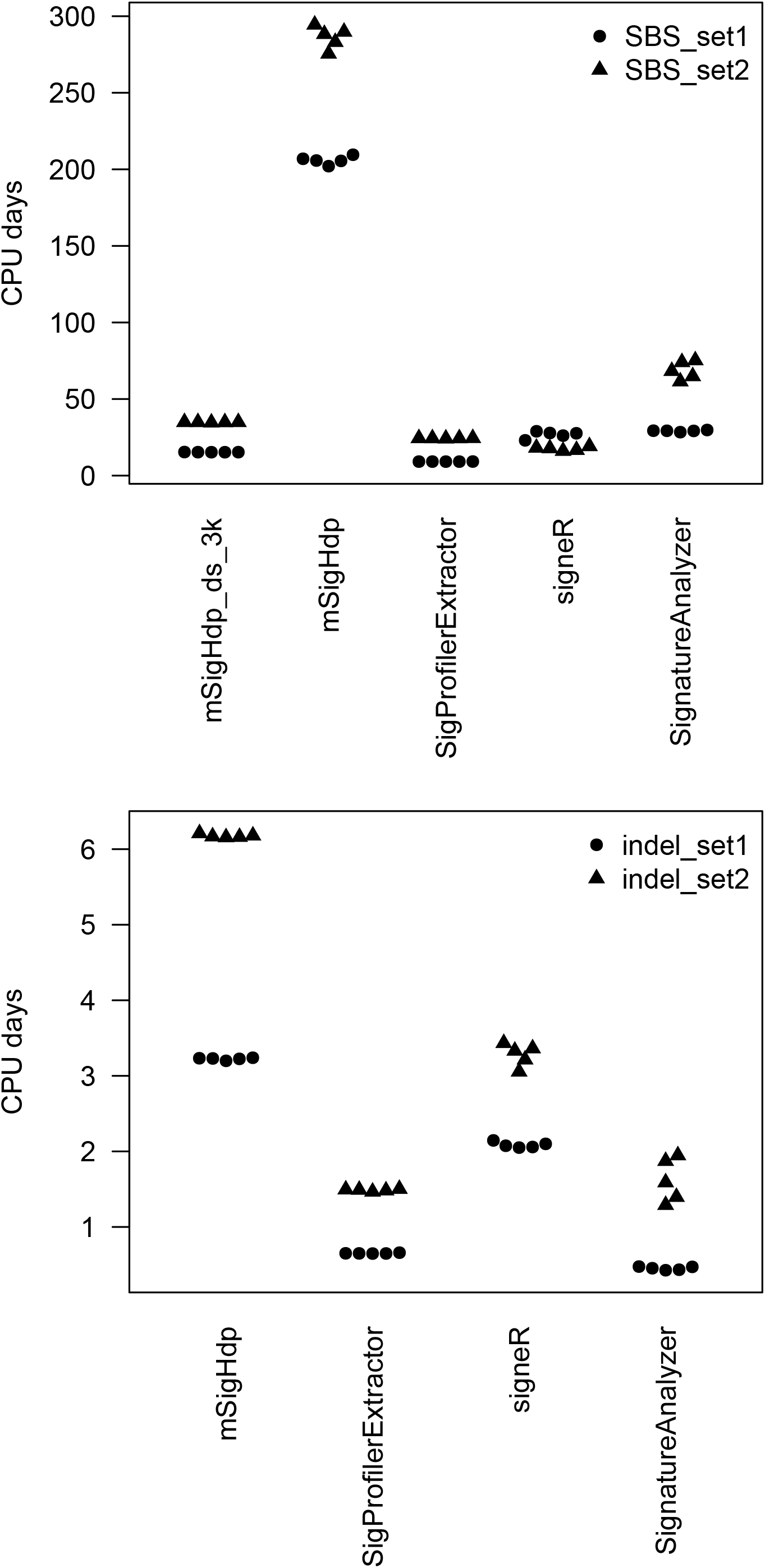
CPU usage for 5 approaches to mutational signature discovery. mSigHdp_ds_3k denotes mSigHdp with a downsampling threshold of 3,000. The y-axis indicates total CPU days used over all subprocesses, including both user and system times. So, for example, for mSigHdp, for which we ran 20 simultaneous Gibbs sampling chains for each run, the “elapsed” CPU days would be approximately 1/20^th^ of the total CPU days. Times were measured by the R system.time function or the Python times function. Except as noted in Table 3, measurements were on AMD EPYC 7763 CPUs with a base clock of 2.45 GHz.

**Table 3.**
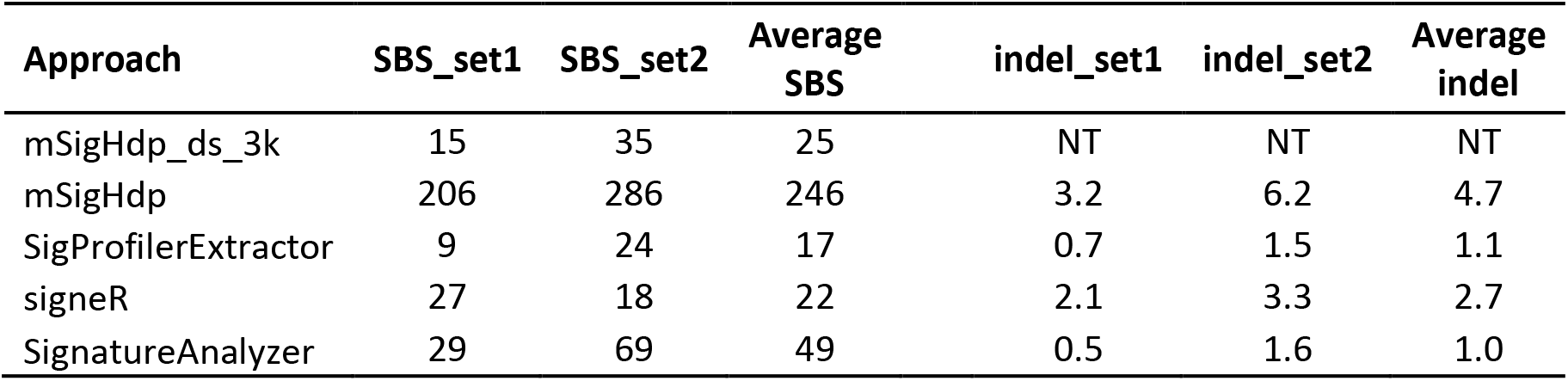
Mean CPU hours needed for 5 approaches to mutational signature discovery. Shown are means over all 5 random seeds. NT: not tested, because mSigHdp with downsampling is not recommended for indel data. mSigHdp_ds_3k denotes mSigHdp with a downsampling threshold of 3,000. Please see Supplementary Table S5 and Figure 7 for more information. We have checked that for signeR the higher CPU usage for SBS_set1 than for SBS_set2 is correct. Measurements were on AMD EPYC 7763 CPUs with a base clock of 2.45 GHz, except for the analysis of SBS data by mSigHdp without downsampling. Because of resource limits on the machines running EPYC 7763 CPUs, for this we used AMD EPYC 7742 CPUs with a base clock of 2.25 GHz.

## Discussion

Here, we have described mSigHdp, software that uses hierarchical Dirichlet process mixture models to discover mutational signatures. We assessed the ability of mSigHdp with and without downsampling to discover mutational signatures in synthetic SBS and indel mutation data, and we compared this to the abilities of 4 other programs. While there has been some systematic benchmarking of mutational signature discovery methods on SBS data (1,6,22–24) we are unaware of previous benchmarking efforts on indel data. Based on an analysis of the reconstruction of mutational spectra in real tumors using known signatures, we believe that the amount of resampling noise in the synthetic SBS and indel mutation data sets used here more closely resembles variability in real data than some previously used benchmarking data (Supplementary Figures S1, S2). Thus, the synthetic mutation data sets here, and notably the indel data set, are resources for developers of mutational-signature discovery methods.

In addition to discovering mutational signatures in sets of mutational spectra, which is the task that mSigHdp and the other approaches tested here carry out, there are other tasks of interest in mutational signature analysis. However, we do not believe mSigHdp or NMF-based mutational signature discovery approaches can be adapted for these other tasks. One of these tasks is the estimation of which already-known mutational signatures have contributed to a given spectrum or (small) set of spectra. This is a hard problem for several reasons. One of these is that if one considers all the many known signatures, there are multiple possible ways to reconstruct the given spectrum, all generating reconstructions that are indistinguishably good. Limiting the set of signatures to those previously observed in a particular tumor type helps but is not sufficient. Multiple programs attempt this task (25–33), but, to our knowledge, there has been no systematic assessment or comparison of their performance. Another task of interest in mutational signature analysis is determining whether a particular signature is present in a tumor. This question has been explored much less, with HRDetect for predicting BRCA1 and BRCA2 deficiency based on mutational signatures being the most prominent example (34). In addition, Ng et al. (35) proposed a generic statistical “signature presence test”(https://github.com/steverozen/mSigAct) and applied it to a question in cancer epidemiology.

The results presented here point to practical suggestions for mutational-signature discovery, especially now that many mutational signatures are well established (1)(https://cancer.sanger.ac.uk/signatures/). A common application of mSigHdp and the other programs benchmarked here would be *de novo* analysis of ≥ 100 mutational spectra to understand what signatures are present and discover possible new signatures. As a practical matter, it seems unlikely that any mutational discovery approach, can discover more than 50 signatures from a single extraction. Currently there are 79 SBS signatures in COSMIC (https://cancer.sanger.ac.uk/signatures/, v3.3, including 19 likely experimental or analytical artifacts). However, these 79 signatures were not discovered in a single analysis of a single large set of 23,000 mutational spectra. Instead, they were discovered by analyses over multiple subsets of 23,000 spectra, including subsets consisting of single cancer types or of tumors heavily exposed to particular mutagens. This is described in detail in Supplementary Note 2 of Alexandrov et al. (1).

Signature discovery is not a lightweight or purely algorithmic endeavor. Indeed, in practice, signature discovery has depended on analyses by multiple programs run on a variety of subsets of the input data and on human judgment drawing on many sources (1). Discovering signatures using different mutation classifications, such as SBSs segregated by translational strand and single-base mutations in the context of the 2 preceding and 2 following bases is also useful, and mSigHdp can do these analyses. Clearly, it would be wise to do several independent analyses, since all methods studied here discovered slightly different sets of signatures when run with different starting random seeds (Figures 5, 6). In any case, one must consider the totality of the evidence, including known signatures, to decide which potentially novel discovered signatures are true positives. Supplementary Note 2 of Alexandrov et al. (1) presents a more extensive discussion of the criteria that are useful for this decision.

In summary, we evaluated the ability of mSigHdp with and without downsampling and of 4 other programs to discover mutational signatures in synthetic SBS and indel mutation data. In the SBS mutation data, mSigHdp with a downsampling threshold of 3,000 was substantially better able to discover signatures than the other programs, and in the indel data, mSigHdp without downsampling was substantially better able to discover signatures (Figures 5, 6, Tables 1, 2). mSigHdp was better able to discover rare SBS signatures, which may be an advantage since most common signatures have probably already been discovered (Supplementary Table S7, Supplementary Figure S10). The results of benchmarking on realistic data here indicate that mSigHdp is an advance for discovering SBS and indel mutational signatures, and especially for discovering rare signatures.

## Acknowledgments

We thank the ICGC/TCGA Pan-Cancer Analysis of Whole Genomes Network for making the mutational spectra, exposures, and signatures available.

