## Supplementary Figures for "mSigHdp: hierarchical Dirichlet process mixture modeling for mutational signature discovery"

[†] Denotes equal contribution.

24 October 2022

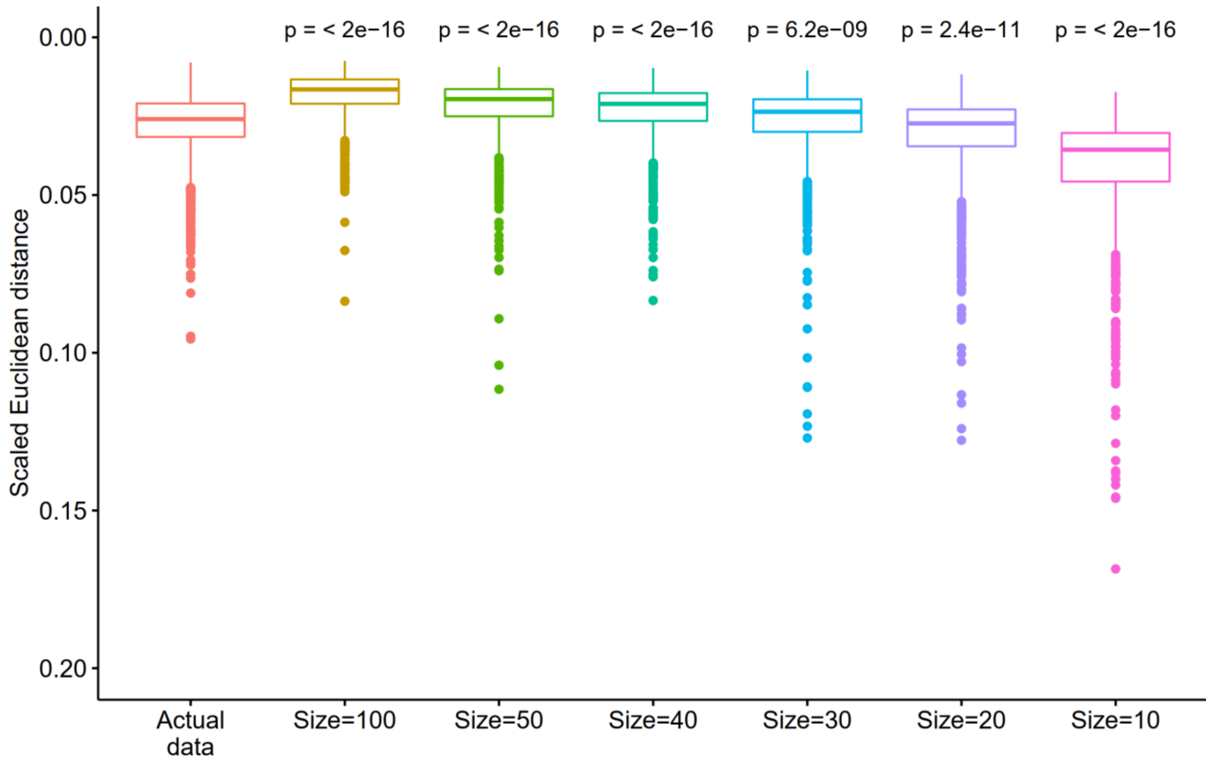


**Supplementary Figure S1 Selecting resampling noise for generating realistic synthetic single-base-substitution (SBS) mutation data across all 18 cancer types.**

We selected the negative-binomial size parameter (indicated as "Size" in the plot) based on the scaled Euclidean distances (Euclidean distance divided by total mutation count) as follows. For the actual data, we used the distance between the actual spectra and the reconstruction of the spectra from mutational signatures and exposures. For synthetic data with a given size parameter, we used the distance between the synthetic spectra with and without resampling noise with the size parameter. Horizontal lines in the boxplots indicate the 1$^{st}$ quartile, median, and 3$^{rd}$ quartile. The ends of the solid lines are at the 1$^{st}$ quartile – 1.5×(interquartile range) and at the 3$^{rd}$ quartile + 1.5×(interquartile range). We selected realistic synthetic data with size = 30 as having the most similar distribution of scaled Euclidean distances compared to the actual data. P values by 2-sided Wilcoxon rank-sum tests versus "Actual data".

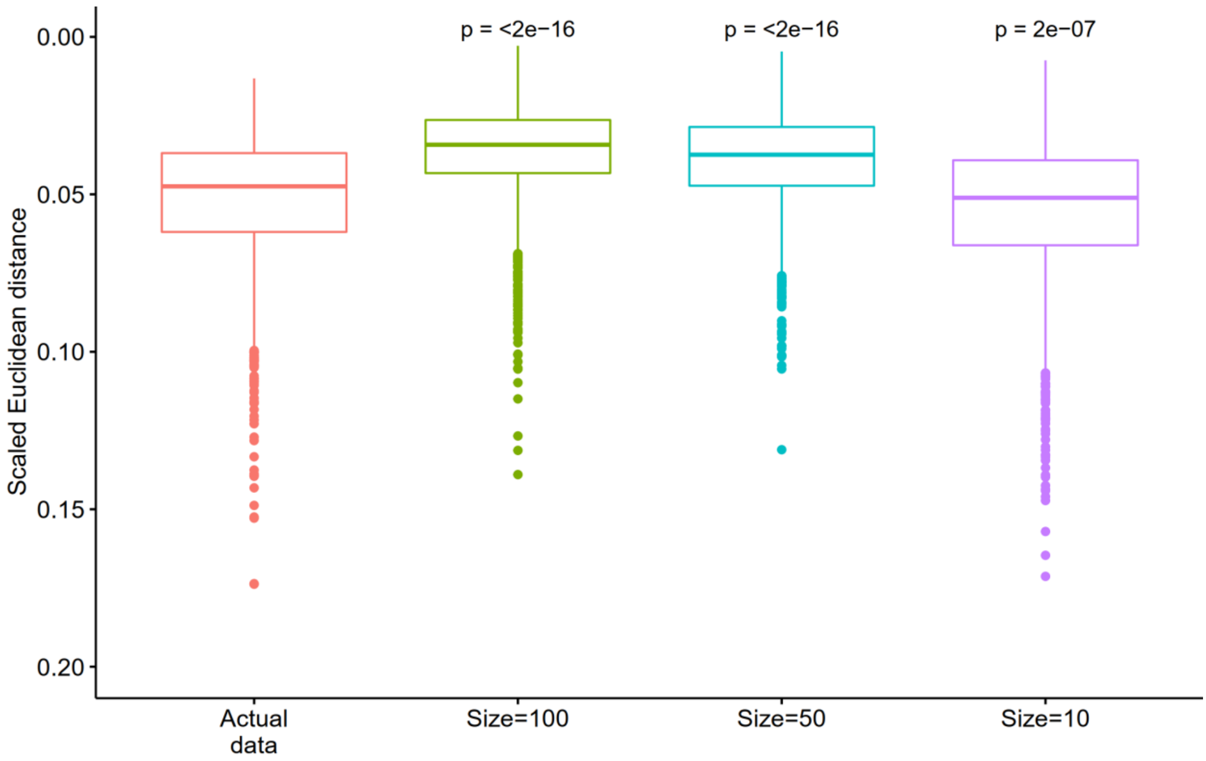


**Supplementary Figure S2 Selecting resampling noise for generating realistic synthetic indel mutation data across all 18 cancer types.**

We selected the negative-binomial size parameter (indicated as "Size" in the plot) based on the scaled Euclidean distances (Euclidean distance divided by total mutation count) as follows. For the actual data, we used the distance between the actual spectra and the reconstruction of the spectra from mutational signatures and exposures. For synthetic data with a given size parameter, we used the distance between the synthetic spectra with and without resampling noise based on the size parameter. Horizontal lines in the boxplots indicate the 1$^{st}$ quartile, median, and 3$^{rd}$ quartile. The ends of the solid lines are at the 1$^{st}$ quartile – 1.5×(interquartile range) and at the 3$^{rd}$ quartile + 1.5×(interquartile range). We selected realistic synthetic data with size = 10 as having the most similar distribution of scaled Euclidean distances compared to the actual data. P values by 2-sided Wilcoxon rank-sum tests versus "Actual data".

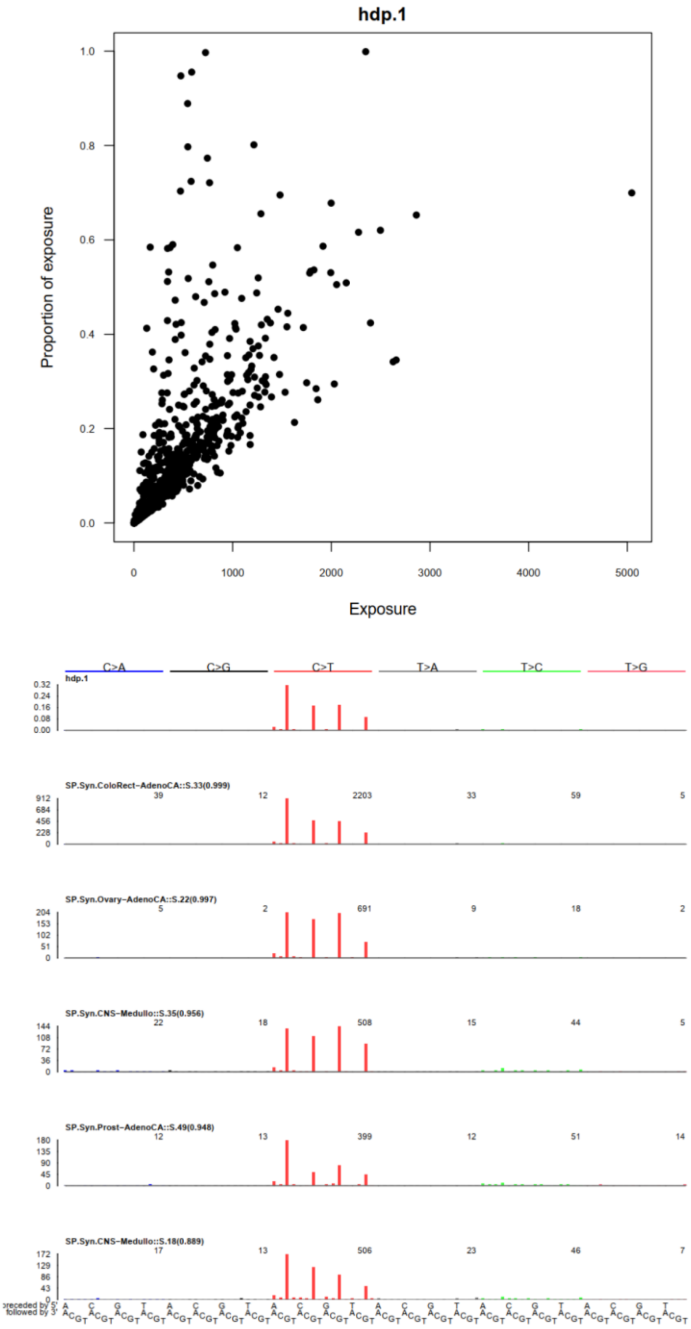


**Supplementary Figure S3 Example mSigHdp diagnostic plots.**

Above: Example of a plot showing the proportion of exposure versus exposure mutation count for a signature discovered by mSigHdp with a downsampling threshold of 3,000. Each dot represents one tumor. Below: Example of a plot showing a signature discovered by mSigHdp (the top plot, labeled hdp.1) and the 5 tumor spectra with the highest proportion of exposure to this signature. The proportion of hdp.1 in each spectrum is indicated in parentheses after the name of the spectrum. From synthetic data SBS_set2, seed 1076753.

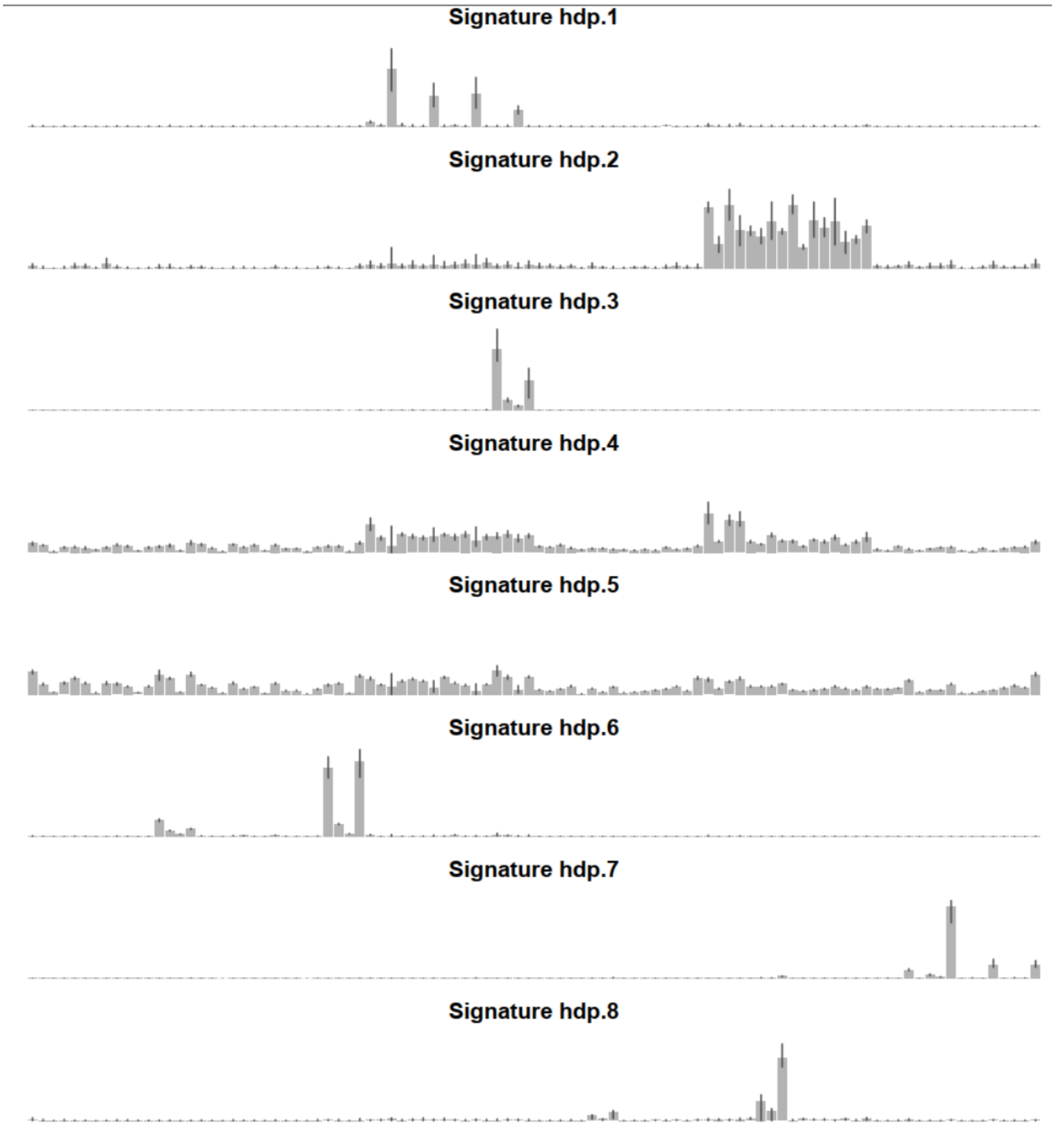


**Supplementary Figure S4 Example mSigHdp diagnostic plots of the profiles of several discovered signatures, with 95% credible intervals of the proportion of mutations of each class.**

The height of each gray bar represents the proportion of mutations in a particular mutation class across all mutations in a group of mutation clusters with similar signature profiles – mutation clusters that represent one mutational signature. See also main text Figure 4. Error bars indicate 95% credible intervals of the height of the associated gray bar, calculated by converting each mutation cluster in a group to its signature profile, and then computing the 95% credible interval of the proportion of each mutation class across all mutation clusters in that group. From mSigHdp with downsampling threshold 3,000 on synthetic data SBS_set2, seed 1076753. This plot was inspired by a similar plot in the original hdp code (https://github.com/nicolaroberts/hdp).

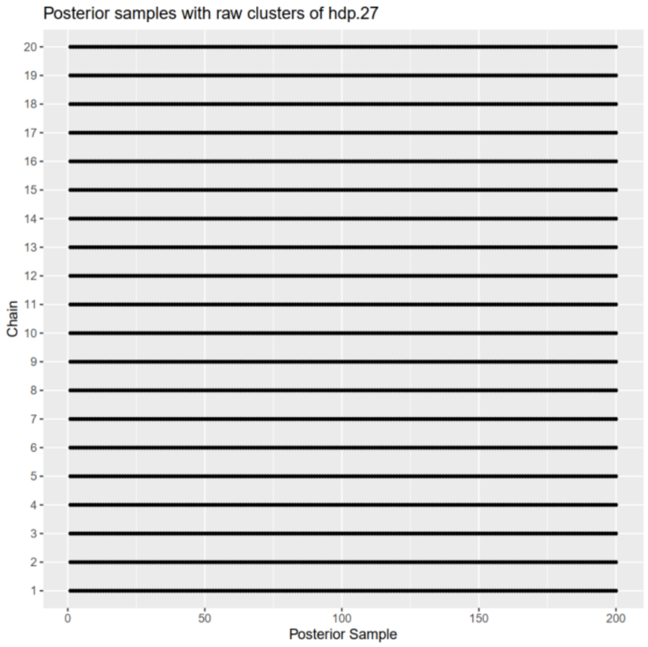


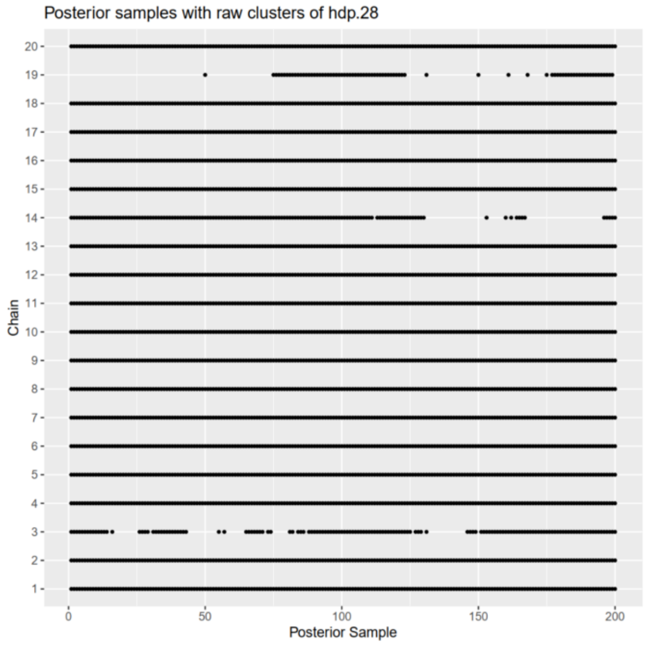


**Supplementary Figure S5 Example mSigHdp diagnostic plots showing which Gibbs samples had mutation clusters that contributed to a particular signature.**

Each plot shows information for one discovered signature in an mSigHdp run that collected 200 posterior samples (indicated as 1 through 200 on the x-axis) on each of the 20 posterior sampling chains (labelled 1 through 20 on the y-axis). There is a dot at each Gibbs sample in which there was a mutation cluster that contributed to a particular discovered mutational signature. Each plot shows real data corresponding to one group of mutation clusters shown in cartoon form in Figure 4B. Above: Clusters of mutations contributing to this signature occurred in all Gibbs samples. Below: Some Gibbs samples in some chains did contribute to this signature. From synthetic data SBS_set2, as analyzed with a downsampling threshold of 3,000 and random seed 1076753.

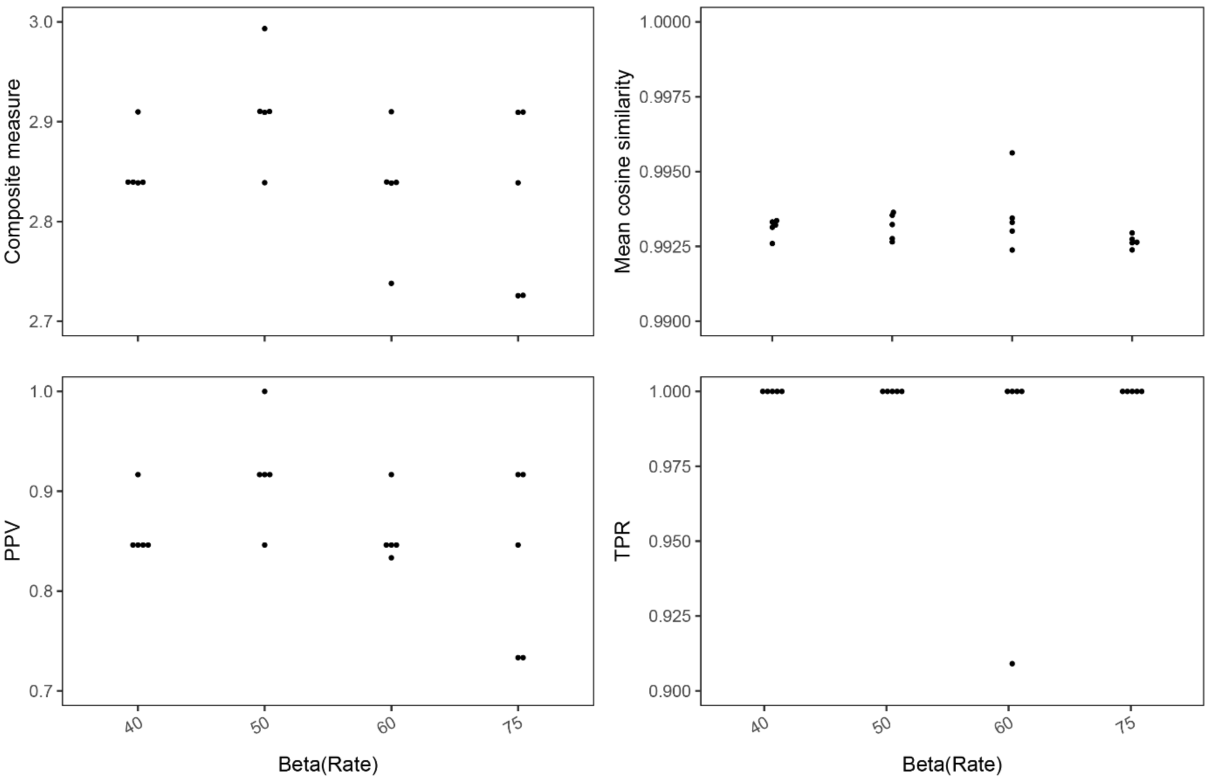


**Supplementary Figure S6 Selecting the prior distribution of Dirichlet process concentration parameters for indel signature discovery.**

Composite measure is the sum of PPV (positive predictive value), TPR (true positive rate), and mean cosine similarity. Each dot indicates a measure for an mSigHdp run on one seed. “Beta (Rate)” refers to the beta parameter of the gamma distribution which serves as the prior distribution of the Dirichlet-process concentration parameters in mSigHdp. We selected beta = 50.

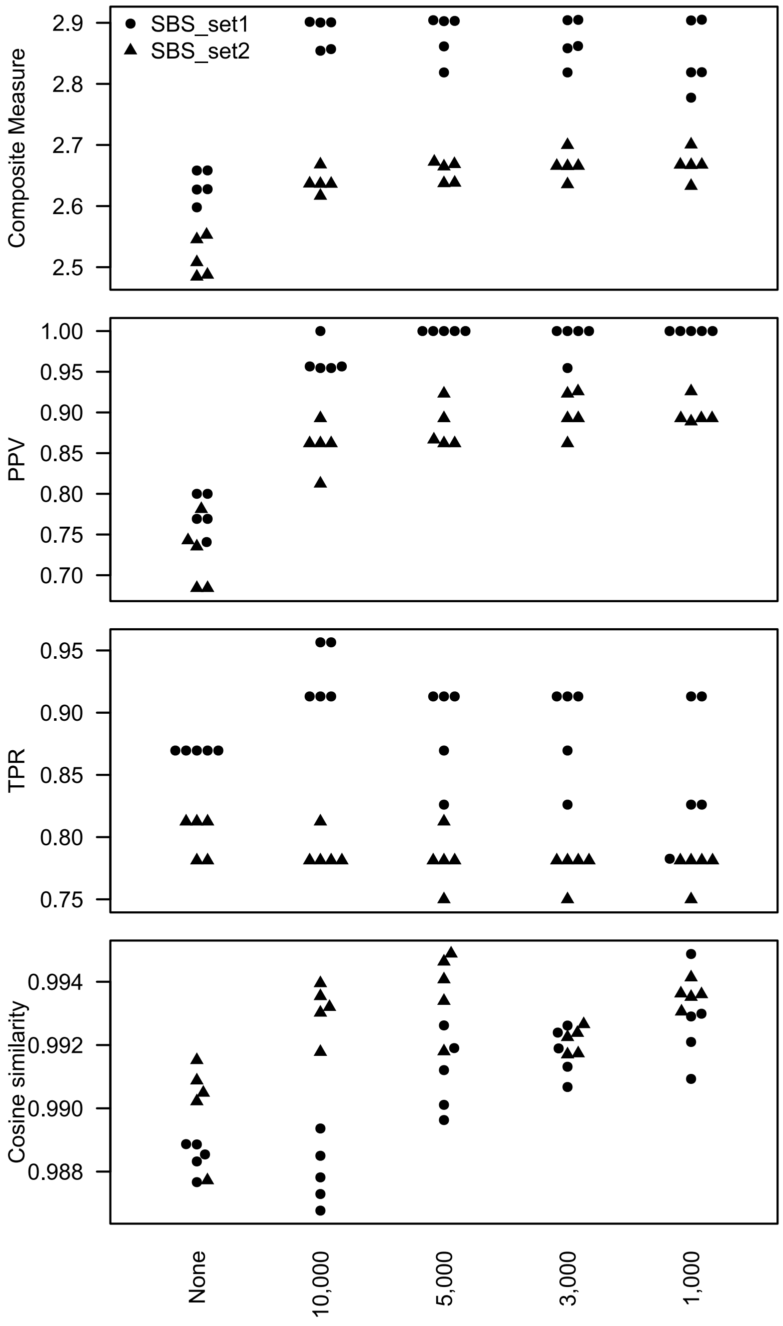


**Supplementary Figure S7 Downsampling SBS mutation data substantially improves mSigHdp's positive predictive value (PPV).**

PPV, positive predictive value; TPR, true positive rate.

We chose a downsampling threshold of 3,000 as having the highest mean Composite Measure across both SBS data sets.

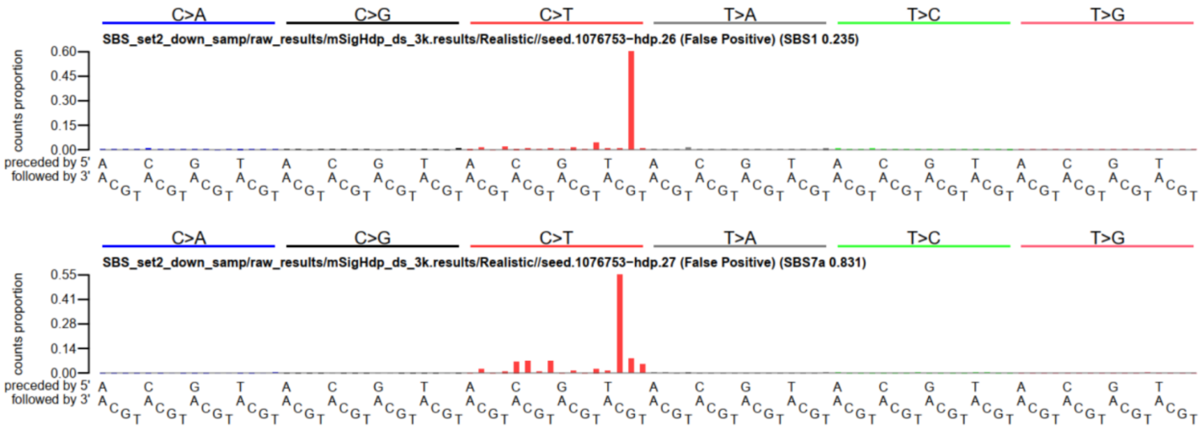


**Supplementary Figure S8 Some false positives discovered by mSigHdp with downsampling consisted of essentially a single mutation class.**

Note that the two plots are different. The top plot shows mutations from TCG → TTG, and the bottom plot shows mutations from TCC → TTC. Other false positives discovered by mSigHdp could not be well reconstructed with false negatives. From SBS_set2, seed 1076753.

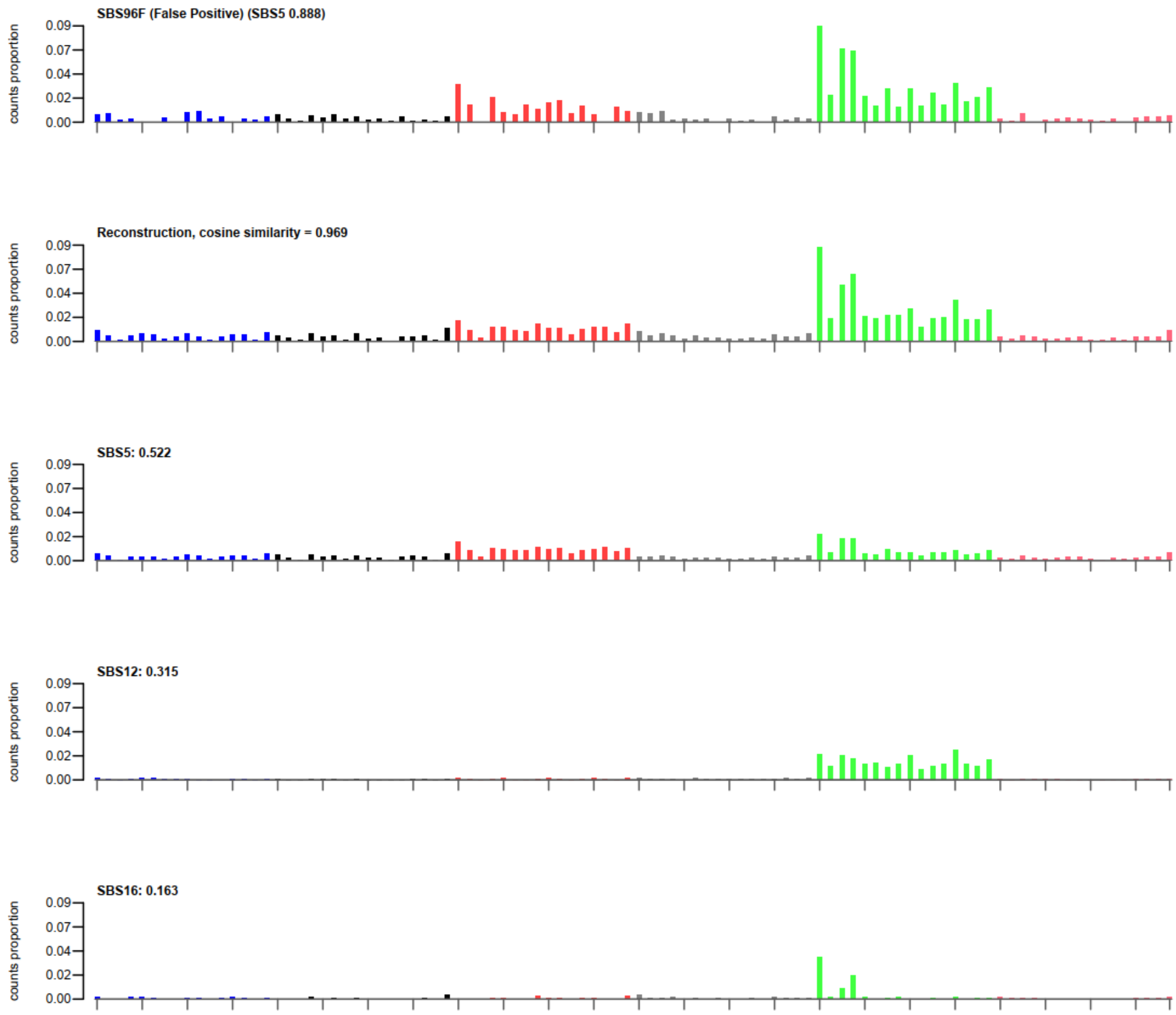


**Supplementary Figure S9 Example of false positive SBS signature discovered by SigProfilerExtractor that could be reconstructed as a combination of 3 false negative signatures.**

The top plot shows the extracted false-positive signature, which had a cosine similarity of 0.889 to signature SBS5. The second plot shows the reconstruction of the extracted false positive signature. This has a cosine similarity of 0.969 to the top plot. The lower three plots show signatures that were added to generate the reconstruction. The proportions used for the reconstruction are shown at the left, namely 0.522 of SBS5, 0.315 of SBS12, and 0.163 of SBS16. The heights of the bars in each of the lower three plots are reduced by the proportion of the corresponding signature to the reconstruction. For example, for SBS5, the height of each bar is reduced by 0.522. Form SBS_set1, seed 200437, mSigHdp with downsampling threshold 3,000.

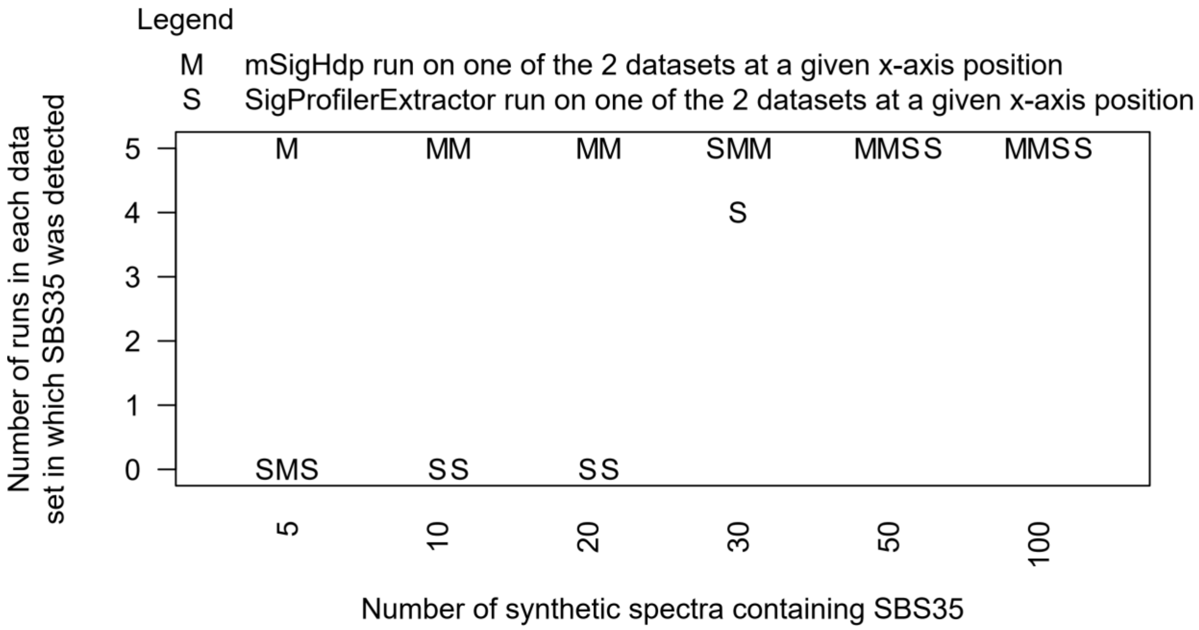


**Supplementary Figure S10 Sensitivity for detecting rare signatures.**

mSigHdp with a downsampling threshold of 3,000 consistently detected SBS35 when it was present in ≥ 10 tumors in 537. We generated the synthetic data from SBS_set1. We first removed tumors already containing SBS35, leaving 537 tumors. Then, for $n \in \{5, 10, 20, 30, 50, 100\}$ we added SBS35 in random amounts (based on the exposures to SBS35 in the "platinum" genomes reported in Alexandrov et al., 2022 [doi: 10.1038/s41586-020-1943-3]) to a random selection of $n$ tumors in 2 replicates. This generated 2 datasets for each $n$ that were independent with respect to the tumors that had SBS35 and with respect to the exposures of these tumors to SBS35. Each program (mSigHdp with downsampling threshold 3,000 and SigProfilerExtractor) was run on 5 random seeds on each of the 2 data sets for a given $n$. Note that for one of the 2 data sets with 30 SBS35-positive tumors, SigProfilerExtractor discovered SBS35 in 4 out of the 5 runs. We chose SBS35 because neither approach discovered it in SBS_set1 or SBS_set2. mSigHdp with a downsampling threshold of 3,000 was able to detect SBS35 at significantly lower prevalence, $p < 7 \times 10^{-3}$ by robust linear regression:

```
MASS::rlm(formula = num.found ~ spike.in.count + Approach,
      data = sbs35_detect)
```

where num.found contains the values on the y-axis of the figure above, spike.in.count contains the values on the x-axis, and Approach is mSigHdp or SigProfilerExtractor. The contents of the sbs35_detect data frame in the function call above are at https://github.com/Rozen-Lab/mSigHdp_sup_files/blob/main/output_for_paper/sup_fig_s10_sbs35_detect_data.csv. The results of the regression are:

| Variable | Value | Std..Error | t.value | p |
|---|---|---|---|---|
| (Intercept) | 3.415 | 0.668 | 5.114 | 4.58E-05 |
| spike.in.count | 0.035 | 0.012 | 2.997 | 6.87E-03 |
| ApproachSigProfilerExtractor | -2.239 | 0.742 | -3.017 | 6.56E-03 |

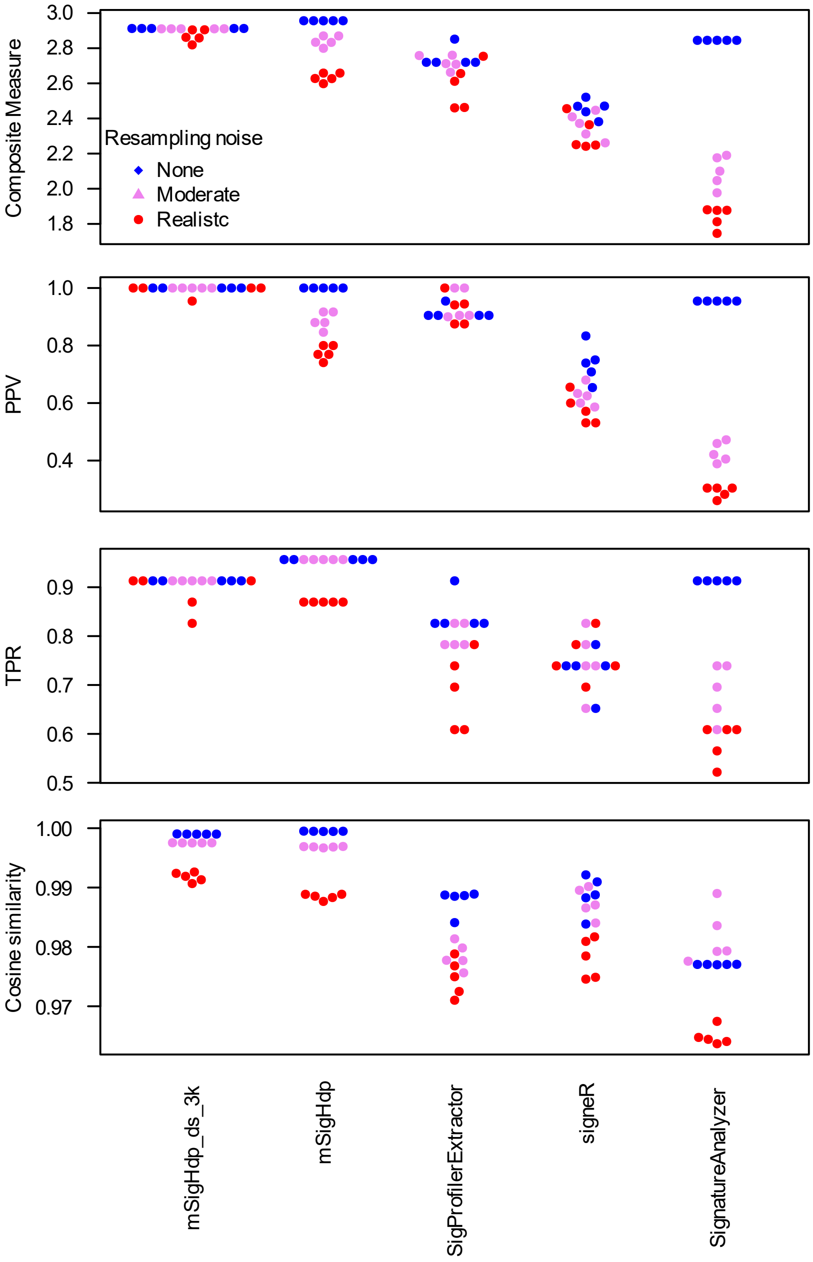


**Supplementary Figure S11 The rankings of mutational-signature-discovery methods were generally unchanged in synthetic SBS data sets with reduced or no resampling noise.**

To gain insight into the sensitivity of methods to different levels of resampling noise in the synthetic input data, in addition to the data with realistic levels of resampling noise (Supplementary Figure S1), we also generated versions of SBS_set1 with "moderate" (less-than-realistic) sampling noise and without resampling noise. For the synthetic data with reduced noise, all programs produced better results. SignatureAnalyzer's results were markedly better for data without resampling noise. mSigHdp_ds_3k denotes mSigHdp with a downsampling threshold of 3,000. For signeR None and Moderate we tested v1.18.1.

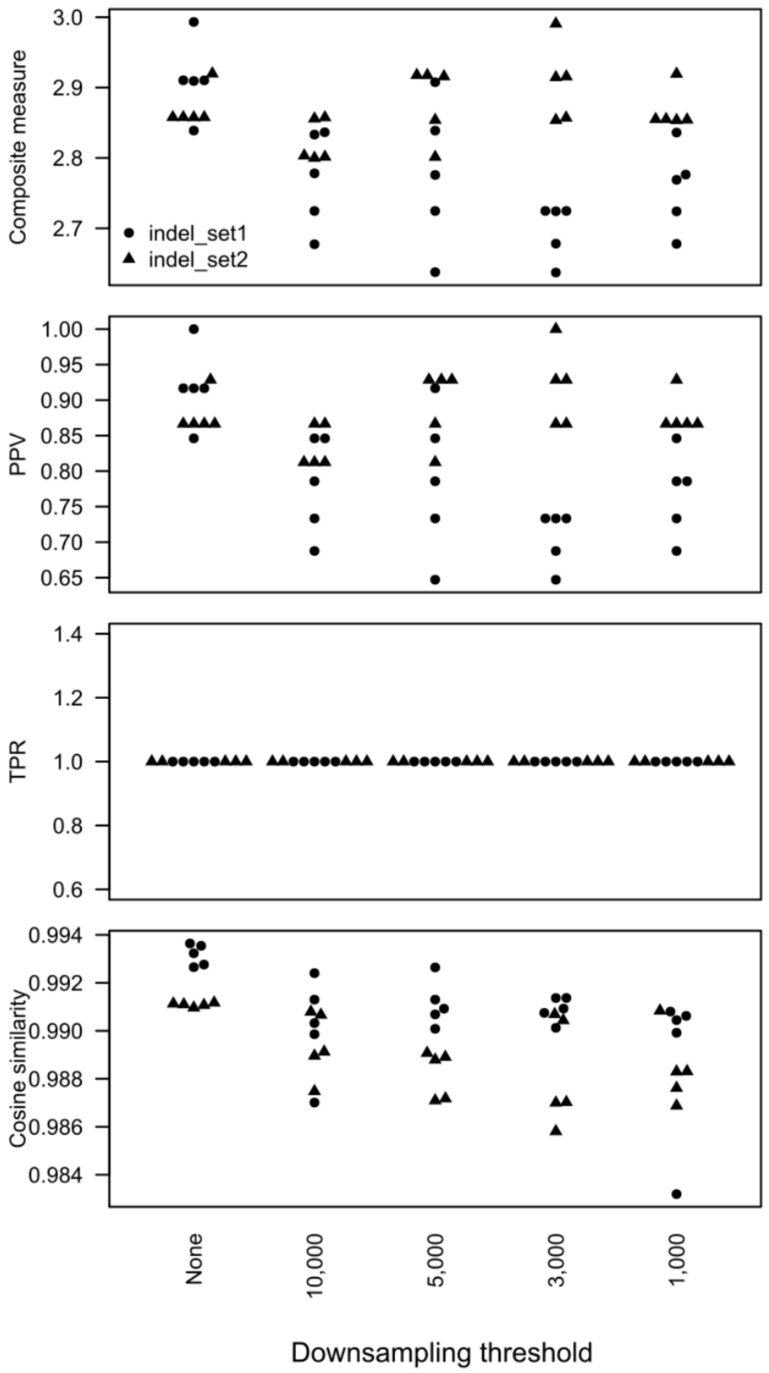


**Supplementary Figure S12 Downsampling of indel data was not useful.**

Note the degraded PPV (positive predictive value) at all downsampling thresholds. TPR: true positive rate.

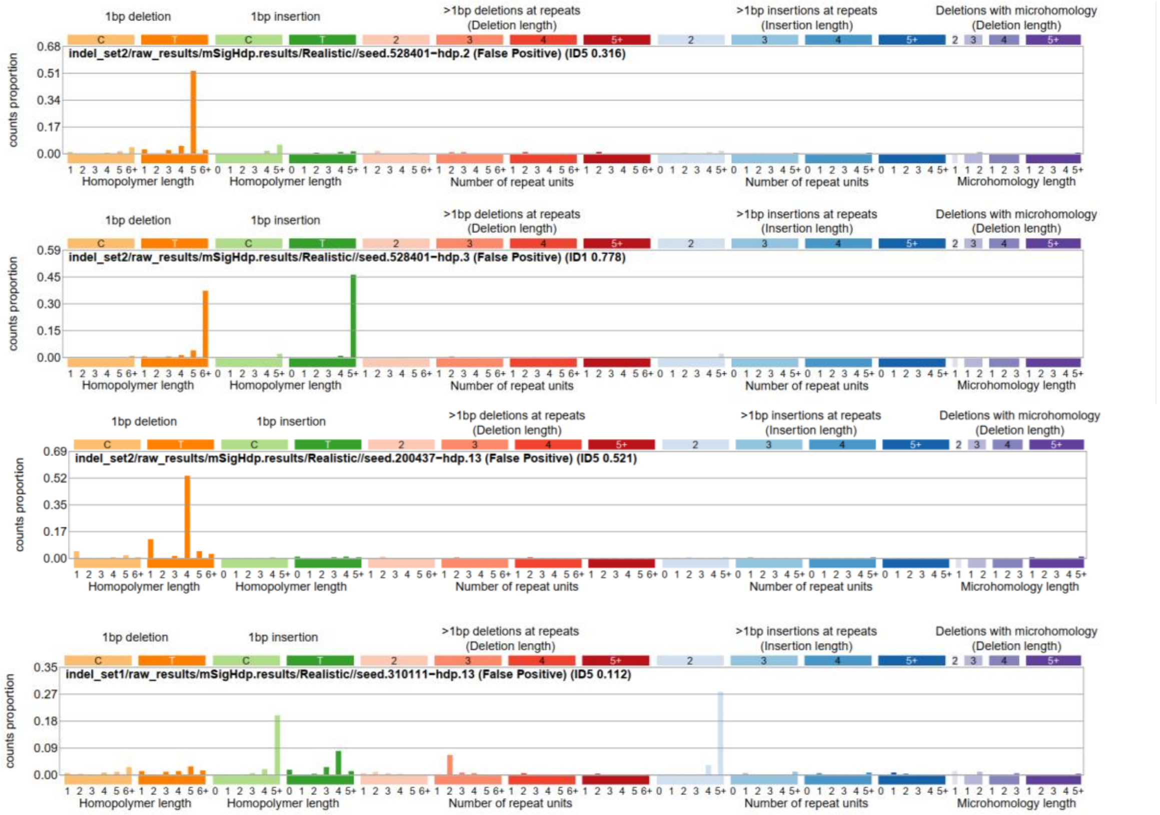


**Supplementary Figure S13 4 mSigHdp indel false positives.**

From top to bottom:

1. A false positive signature that was found in many runs of both indel_set1 and indel_set2.
2. A false positive in many runs of indel_set2.
3. A false positive in a single indel_set2 run.
4. A false positive in a single indel_set1 run.

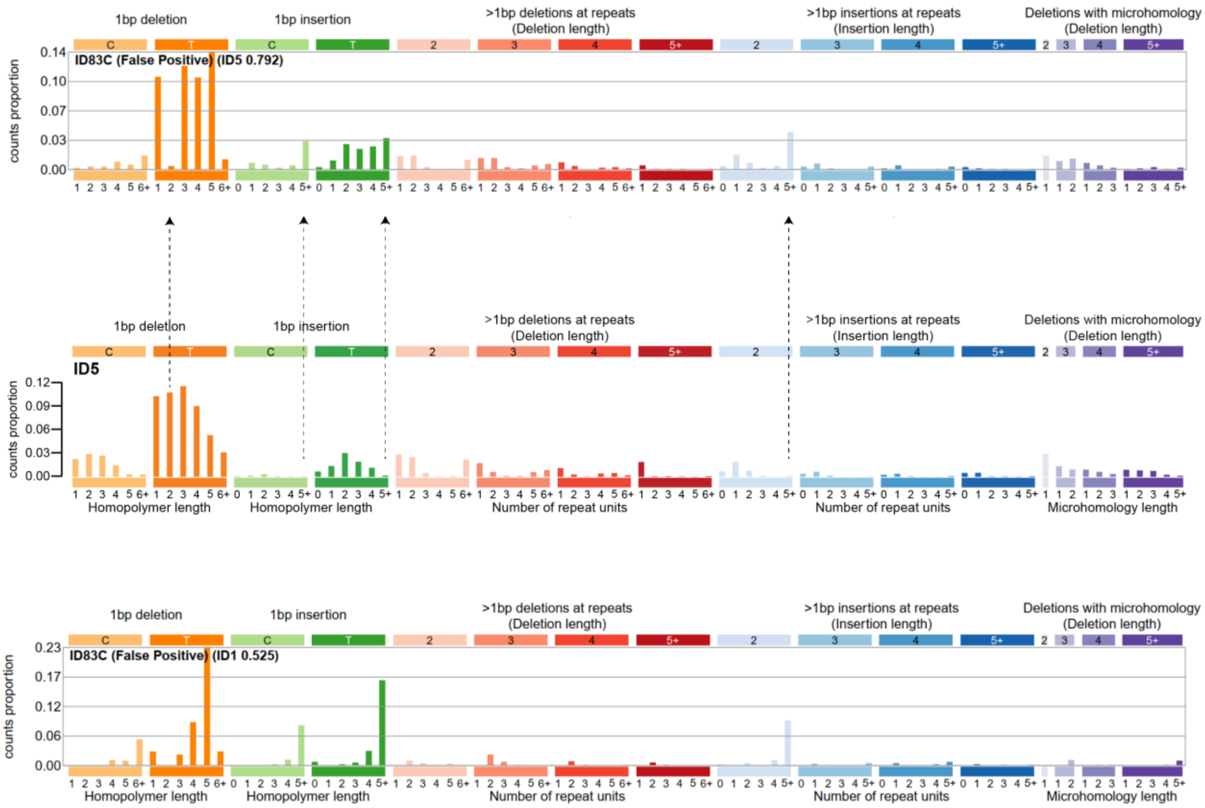


**Supplementary Figure S14 Indel false positive signatures discovered by SigProfilerExtractor.**

Above, a false positive signature resembling ID5 but with one mutation class absent and some additional mutation classes added (absent and added mutation classes indicated by arrows) was discovered in both indel_set1 and indel_set2. The mutational signature ID5 is from https://cancer.sanger.ac.uk/signatures/id/id5/, v3.2. The false positive is from indel_set1, seed 1076753. Below, false positive found in analyses of indel_set2 (seed 1076753).

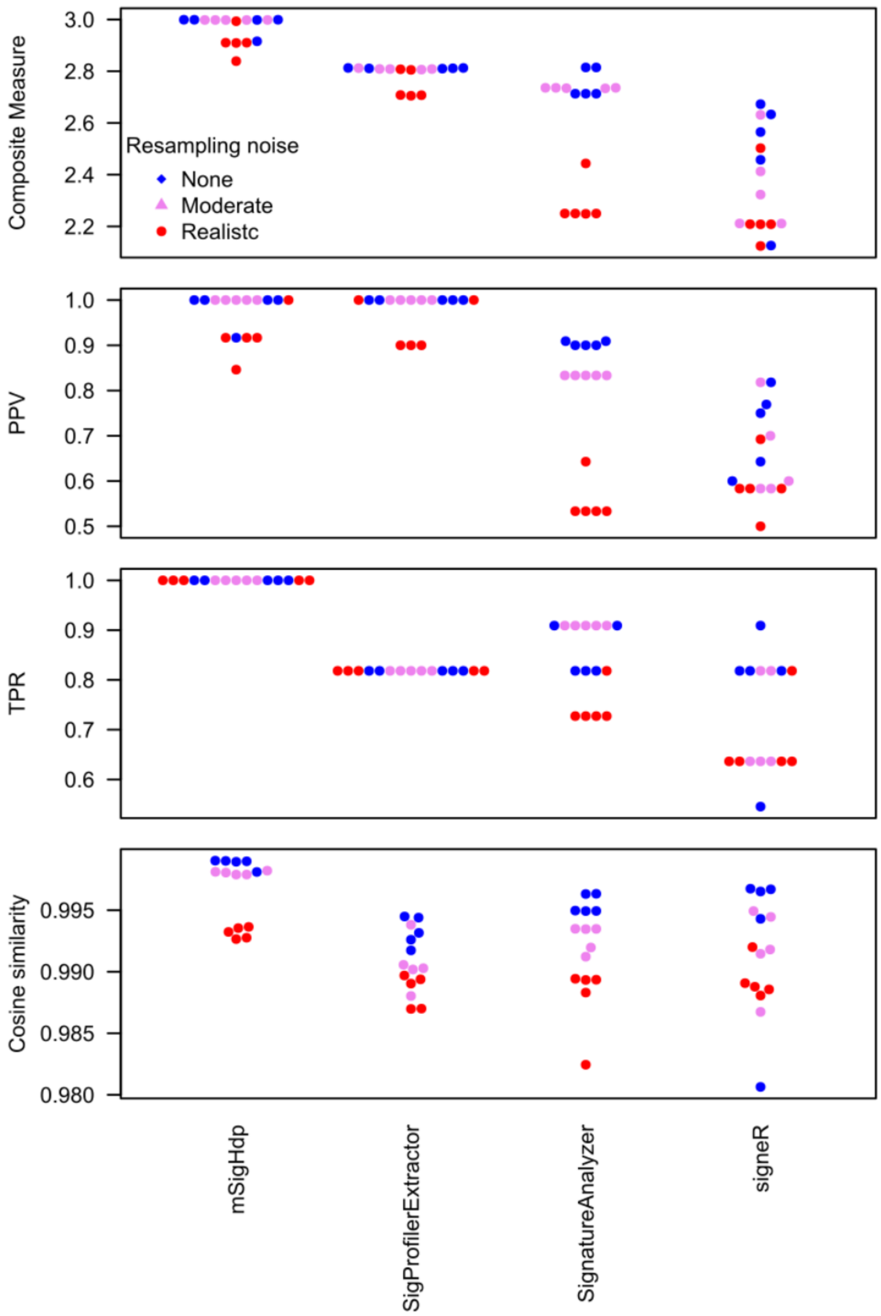


**Supplementary Figure S15 The rankings of mutational-signature-discovery methods were generally unchanged in synthetic indel data sets with reduced or no resampling noise.**

To gain insight into the sensitivity of methods to different levels of resampling noise in the synthetic input data, in addition to the data with realistic levels of resampling noise (Supplementary Figure S2), we also generated versions of indel_set1 with “moderate” (less-than-realistic) sampling noise and without resampling noise. For the synthetic data with reduced noise, all approaches produced better results. SignatureAnalyzer’s results were markedly better for data with less resampling noise.
